# Bioinformatics-Driven Identification and Prioritization of PTSD Targets Based on Published Multi-omic Data

**DOI:** 10.1101/2025.04.07.647615

**Authors:** Mark Zervas, Allyson Gage, Magali Haas

## Abstract

Efforts in recent years to uncover neurobiological mechanisms underlying post-traumatic stress disorder (PTSD) have yielded an expanding candidate pool of targets from genomic and transcriptomic data. However, not all candidates are disease-causing, related to pathological mechanisms, clinically relevant, nor druggable by conventional means. An effective method to systematically identify and prioritize high-confidence, high-impact targets in the central nervous system (CNS) is required to de-risk resource-intensive experimental validation of disease mechanisms and accelerate the development of novel treatments. Here, we describe methods and implementation of a novel 3-phased, biologically rationalized, and quantitative prioritization strategy to identify and rank PTSD-associated targets based on confidence of association to PTSD and estimated CNS-relevant pathogenicity. *Phase 1* was designed to identify and advance targets confidently associated with PTSD through their expression in CNS tissues. Putative targets derived from 29 transcriptomic and genomic analyses of PTSD were evaluated for: 1. Replication in independent cohorts, 2. Observation of differential expression in PTSD CNS tissues, and 3. Demonstration of consistent direction of effect. This strategy resulted in 177 targets that passed criteria for advancement. Phase 1-selected targets were enriched for PTSD relevant traits including irritability, emotional symptoms, and insomnia (FDR <0.05). *Phase 2* advanced targets with additional evidence of association to pathological CNS phenotypes. DisGeNET gene-disease association scores were applied to each Phase 1-selected target to assign a confidence score indicating that a target was associated with CNS-relevant pathology using criteria for moderate or strong evidence of CNS disease association. Phase 2 advanced 55 of the 177 (31.1%) targets. The Phase 2 target pool was enriched for CNS phenotypic abnormalities (FDR<0.05). Finally, *Phase 3* enabled target prioritization by annotating targets with a composite pathogenicity score. Components of the pathogenicity score included metrics derived from drug trial databases, predicted loss-of-function intolerance, and connectivity within a protein-protein interaction network defined by PTSD-associated targets. The resulting 55 targets were ultimately prioritized by the sum of Phase 2 and Phase 3 scores, where top-ranked targets had strong evidence to support both association with PTSD in brain and high pathogenicity estimates in a CNS-relevant context. Biologically, top-ranked targets implicate transmitter systems (GABA, histamine, and estrogen), structural regulation of neurites, and protein homeostasis. Future work will be required to experimentally validate the utility of the high priority PTSD targets we identified as well as to demonstrate the general applicability of this methodology. Ultimately, we anticipate that the three phased approach will enable efficient de-risking of PTSD and other poorly understood CNS disorders.

## INTRODUCTION

Post-Traumatic Stress Disorder (PTSD) is the fourth most common psychiatric condition in the United States, affecting both military and civilian populations, and contributes to a substantial public health burden. PTSD is currently diagnosed based on clinical symptoms, including re-experiencing, avoidance, negative cognitions and mood, and arousal [Diagnostic and Statistical Manual of Mental Disorders, Fifth Edition, Text Revision (DSM-5-TR)]^**1**^. Available therapies exist, but are limited in treating PTSD, including only two FDA approved <u>s</u>elective <u>s</u>erotonin reuptake inhibitors (SSRIs)^**2–4**^, which have limited efficacy, undesirable side effects, and target only the symptoms of PTSD, but not cellular pathology, largely because of the sparsity of specific pathways and mechanisms underlying the disorder. As a result, no additional approvals have been issued for PTSD in over 20 years including a recent FDA complete response letter for MDMA-assisted therapy^**64,65**^. The PTSD drug development deadlock is driven by multiple contributing factors including blinding, the lack of understanding of the biological underpinnings of PTSD, the poor predictive value of preclinical models^**59,60**^ (e.g. Single Prolonged Stress^**61**^), a lack of disease relevant and predictive therapeutic biomarkers, and the challenges of identifying potent molecules within the stringent range of molecular properties necessary for blood-brain barrier (BBB) penetration^**62**^. Thus, there is a need for rational drug development to identify and advance novel PTSD therapeutic targets.

Based on the premise that the expression of PTSD relies on a complex interaction between severity of exposure to traumatic stress and underlying biological and genetic susceptibility, Cohen Veterans Bioscience, a nonprofit biomedical research organization, invested in research to improve the probability of drug discovery and biomarker success. Initial investments focused on evaluating genetic susceptibility for PTSD through the establishment of a PTSD Consortium to lead the first analysis of global <u>g</u>enome <u>w</u>ide <u>a</u>ssociation studies (GWAS)^**5–8**^. This program established that PTSD susceptibility is heritable and highly polygenic^**5**^, which reinforces that PTSD has a biological underpinning. However, there are significant challenges in mapping <u>s</u>ingle <u>n</u>ucleotide <u>p</u>olymorphisms (SNPs) identified in GWAS to definitive, select genes^**9–11**^, which limits the identification of causal pathways and therapeutic targets. To understand how diverse genetic perturbations converge onto functional pathways, CVB funded a project that resulted in the development of a PPI network encoded by genes strongly associated with PTSD based on SNPs identified by the PTSD Psychiatric Genomics Consortium^**57**^.

Alternative approaches for target identification include conducting bulk (tissue) or cell-type specific (single cell, sc) RNA <u>seq</u>uencing (RNAseq) studies, which have been applied both broadly to developmental disorders^**12**^ and specifically to PTSD^**13–16**^. Unsurprisingly, a number of brain regions are implicated in PTSD including the prefrontal cortex (PFC) and the amygdala. Accordingly, studies focused on human RNAseq (aka transcriptomics) have been conducted on specific regions derived from postmortem CNS tissue (e.g. PFC subdomains) and from human iPSC-derived <u>exc</u>itatory <u>neu</u>rons (iPSC-Neu^Exc^)^**13–16**^. Importantly, these studies have provided lists containing transcripts that may play a role in PTSD. The utility of RNAseq has been demonstrated for recent drug discovery efforts, yielding actionable targets, in particular in the context of oncology drug development^**17–19**^. With the ultimate goal of facilitating novel therapeutic discovery efforts to ameliorate PTSD, we set out to identify and prioritize PTSD-relevant targets that would further our understanding of neuronal phenotypes underpinning PTSD. Knowing that we could not confidently associate GWAS-identified SNPs with definitive causal genes, we took advantage of the publicly available transcript-based data supplemented with published GWAS^**13–16**^ to identify target genes with increased confidence.

Notably, transcriptomic studies typically identify large numbers of transcripts that are either up-regulated or down-regulated. Thus, identifying and prioritizing transcripts using large transcriptomic data sets is a challenge due to the number of possible targets and their specific relevance to underlying cellular pathology. Therefore, we developed a novel 3-phase quantitative prioritization strategy to identify and rank PTSD-associated transcripts derived from publicly available sources^**13–16**^ and using the principle of consilience we ultimately prioritized and selected 20 ‘index’ transcripts and their related proteins to further interrogate and generate PTSD <u>p</u>rotein-<u>p</u>rotein interaction (PPI) networks. Our central hypothesis was this three phased approach will advance PTSD disease understanding and therapeutic development by contributing to knowledge of PTSD neuronal phenotypes and disease signatures. Ideally PPI networks, derived from our selected index proteins, would be functionally validated using a series of experimental assays, as part of future research efforts. Finally, we believe our approach, which we applied to PTSD as a practical example, will be applicable to other complex disorders to aid in target identification.

## METHODS

The following describes a novel, rationalized, and systematic three-phased bioinformatics prioritization strategy (**Fig. 1**) to (i) Nominate independently replicated PTSD-associated targets, (ii) Determine their observed differential expression in PTSD brain tissues, and (iii) Characterize evidence to support CNS-relevant pathogenicity for eventual advancement to functional assays using induced <u>exc</u>itatory <u>neu</u>rons (iNeu^Exc^)^**25**^, which capture cellular disease phenotypes and are distinct from iPSC-Neu^Exc^, which eliminate phenotypes due to reprogramming to a ground state. We utilized a number of publicly available tools to facilitate our efforts: STRING^**20,21**^ is a free PPI database that captures physical and functional interactions, parsed on species (e.g. human), which can be viewed in STRING, but can also be imported, visualized, and further analyzed via a plugin module with Cytoscape^**22–24**^. As protein-protein associations may not necessarily indicated physical interactions, we took an additional (first neighbors) step in Cytoscape to build on this core network and expand the PTSD protein interaction network emphasizing likely physical interactions. The three phases are:

**Fig. 1.**
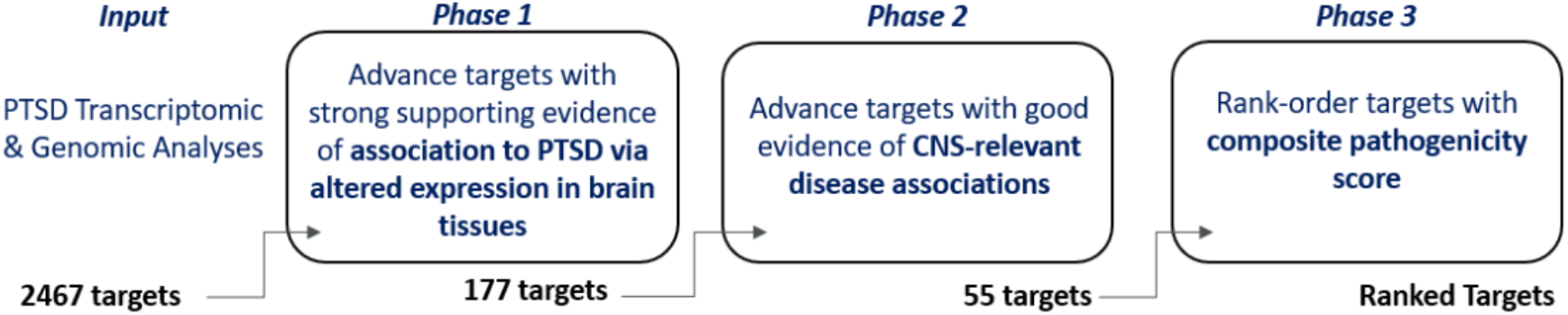
Overview of PTSD Brain Target Prioritization Strategy. Transcriptomic and genomic datasets were assessed to identify targets in brain tissues/iPSC-Exc^Neu^ that are confidently associated with PTSD (Phase 1, yielding n=177 targets) and associated with other CNS phenotypes using DisGeNET (Phase 2, yielding n=55 targets), and are likely to be pathogenic based on a three-component pathogenicity score (Phase 3) ultimately leading to a cohort of top ranked PTSD targets (n=20).

**Phase 1**. All data and calculations related to Phase 1 are provided in **Supplemental Table 1**. Candidate PTSD-associated transcriptomic targets and their direction of regulated expression were extracted from 10 studies, encompassing 29 analyses. Across each study, potential targets were annotated as to whether they represented (*i*) Direct observation of differential transcript expression in brain tissues or in iPSC-Neu^Exc^ from PTSD cases versus controls^**13–16**^, (*ii*) Imputation of differential transcript or protein expression in brain tissues in PTSD cases versus controls^**13,26–28**^, and (*iii*) Inference of PTSD-associated gene through GWAS^**5–8,29**^. The details of each analysis are in **Supplemental Table 1**. The four criteria listed below reduced presumptive targets (2467) to generate an actionable list:

1. Directly observed to be differentially expressed in PTSD brain tissues or iPSC-Neu^Exc^: n=4 cortical tissues, n=1 subcortical tissue (amygdala), and n=1 cell type (excitatory neuron) are represented.
2. Observed in independent cohorts: n=6 fully independent cohorts represented in the data set.
3. Exhibited consistent (>50%) direction of difference relative to controls across all analysis in which it was identified.
4. At least one of the following:

- One or more *additional* observations of differential expression in PTSD brain tissues or neurons.
- Supported by imputation of differential expression in brain tissues from PTSD GWAS data (for review of these methods, see Li et al., 2021).^**30**^
- Implicated by a genome-wide significant SNP from a publicly available PTSD GWAS.

Ultimately, the utility of Phase 1 was to generate an actionable list of targets (n=177), which was defined as an economically viable set of transcripts/proteins that may be experimentally evaluated by most academic or industry labs while still revealing molecular signatures that provide insight to underlying pathogenic mechanisms.

**Phase 2**. All data and calculations related to Phase 2 are provided in **Supplemental Table 2**. DisGeNET (www.disgenet.org) <u>g</u>ene <u>d</u>isease <u>a</u>ssociation (GDA) scores were obtained from DisGeNET and used to establish orthogonal evidence for target relevance in a CNS disease context. DisGeNET GDA scores reflect the strength of evidence for a gene-disease linkage across multiple databases (**Fig. S2**), have previously been used successfully in CNS target prioritization, and include bioinformatic/computational efforts in support of neurodegeneration and Parkinson’s Disease^**31,32**^. DisGeNET has also validated prediction efforts using either *in vitro* or *in vivo* models of Alzheimer’s Disease^**33**^ and ADHD^**34**^. In a drug development context, a retrospective analysis found that strong DisGeNET GDA scores outclassed all other parameters in increasing chances of success from phase 1 to launch^**35**^. Thus, targets with GDA scores <0.3 were deprioritized, which left 55 targets to be advanced to Phase 3.

**Phase 3**. Targets (n=55) advanced to Phase 3 were associated to PTSD through altered brain tissue or neuron expression (Phase 1) and have linkages to CNS-relevant phenotypes (Phase 2). The goal of Phase 3 was to prioritize targets allowing for efficient use of resources during future validation experiments.

Therefore, we integrated additional metrics that estimated a target’s pathogenicity and generated a composite pathogenicity score, which was ultimately combined with the Phase 2 GDA score and yielded 20 top tier PTSD targets. All Phase 3 data and calculations are provided in **Supplemental Table 3**. To benchmark effects of Phase 1 on the candidate target pool, enrichment analysis using the Monarch human phenotype-genotype database^**36**^ was performed with the target pool before (n=2467) and after (n=177) Phase 1-selection, using the whole genome for statistical background (analysis tool is available through www.string-db.org).

## RESULTS

### PTSD Target Identification

<u>The goal of phase 1</u> was to identify and advance targets with strong supporting evidence of association with PTSD via altered expression in brain tissues (**Fig. 1**). We combed through 10 studies and identified 2467 potential targets. Of these, 177 targets met our four criteria, listed above, and were advanced to Phase 2 (**Supplemental Table 4, Fig. S2**). While candidate targets prior to selection showed some (minor) enrichment for relevant phenotypes such as panic attack (**Fig. 2A**), there was a notable emergence (5-fold increase) of multiple psychiatric and CNS disease phenotypes that resulted *following* Phase 1 selection (**Fig. 2B**). All terms derived from Phase I were significant at FDR <0.05. Note that “Strength” on the x-axis represents Effect Size, which is defined in the Fig. 2 legend, and reflects enrichment of specific phenotypes. These results indicate that advancing putative targets on the basis of a reproducible RNAseq signal derived from PTSD tissues/ iPSC-Exc^Neu^ enriched the target pool with candidates implicated in neuronal/CNS phenotypes, which we believe have relevance to PTSD. Subsequently, targets that survived Phase 1 were advanced to a second enrichment step to further refine the target list.

**Fig 2.**
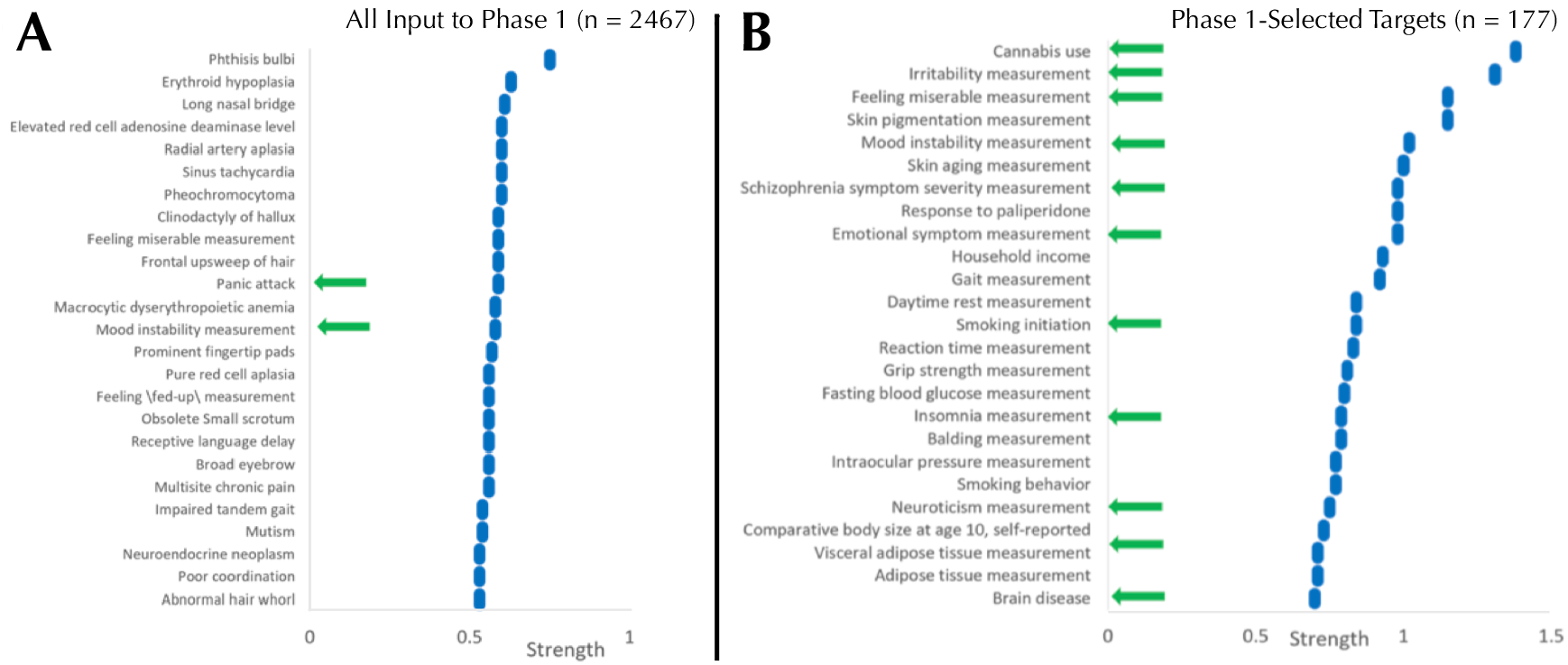
Phase-1 Selected Targets were Enriched for CNS Phenotypes. Panels represent the top 25 terms resulting from Monarch human phenotype enrichment analysis (via STRING) using all PTSD candidate targets (n=2467) identified in source publications (**A**) and phase 1-selected targets (n=177, **B**). Targets selected in Phase 1, reflecting confidence of association to PTSD in brain tissues, were enriched for multiple PTSD-relevant phenotypes, including irritability, emotional symptom measurement, and insomnia (**B**). All analyses were conducted using the whole genome as the background. “Strength” on the x-axis represents <u>Effect Size</u>, which equals log10 [(Observed number of proteins in the analyzed network annotated to phenotype ‘X’:Total number of all proteins annotated to phenotype ‘X’):(Expected number of proteins annotated to Phenotype ‘X’ in a random network of the same size)]. For perspective, an effect size of 0.3 would be equivalent to a 2-fold enrichment.

<u>The goal of phase</u> 2 was to advance Phase 1-nominated PTSD-associated targets that have orthogonal evidence of association to CNS pathological (disease) phenotypes (**Fig. 1**). CNS-relevant GDA scores were defined by disease terms containing “*nervous system disease*”, “*mental disorder*”, or “*behavioral mechanisms*” and were first annotated to their respective Phase 1 PTSD relevant targets (**Fig. 3**). Each target was categorized by its highest GDA score as having “weak” (< 0.3), “moderate” (0.3-0.39), or “strong” (≥ 0.4) effect size. Among the 177 Phase 1 targets, 135 (77.4%) had at least one CNS-relevant GDA of any strength, with 55 (31.07%) possessing at least one moderate or strong CNS-relevant GDA (**Fig. 2 and Fig. 3**). Moderate and strong gene-disease associations, which included linkages to schizophrenia, autism, depression, neurodegeneration (and more) were archived in a PTSD gene-disease linkage map for future reference; this can be found in **Supplemental Table 2**. The 55 targets that had at least one moderate or strong CNS-relevant GDA were advanced from Phase 2: For simplicity, GDA scores were assigned a “Phase 2 Score” of 3 (i.e. with strong GDA, n=34) or “Phase 2 Score” of 2 (i.e. with moderate GDA, n=21) (**Table 1**). To validate that the Phase 2 selection procedure, based on strength of the DisGeNET GDA score, enriched the target list for disease an enrichment analysis was performed using Monarch human phenotype-gene associations^**36**^. Specifically, Phase 2 targets were tested for enrichment against a background of Phase 1 targets, revealing significant enrichment for CNS phenotypic abnormalities (**Fig. 4**).

**Table 1.**
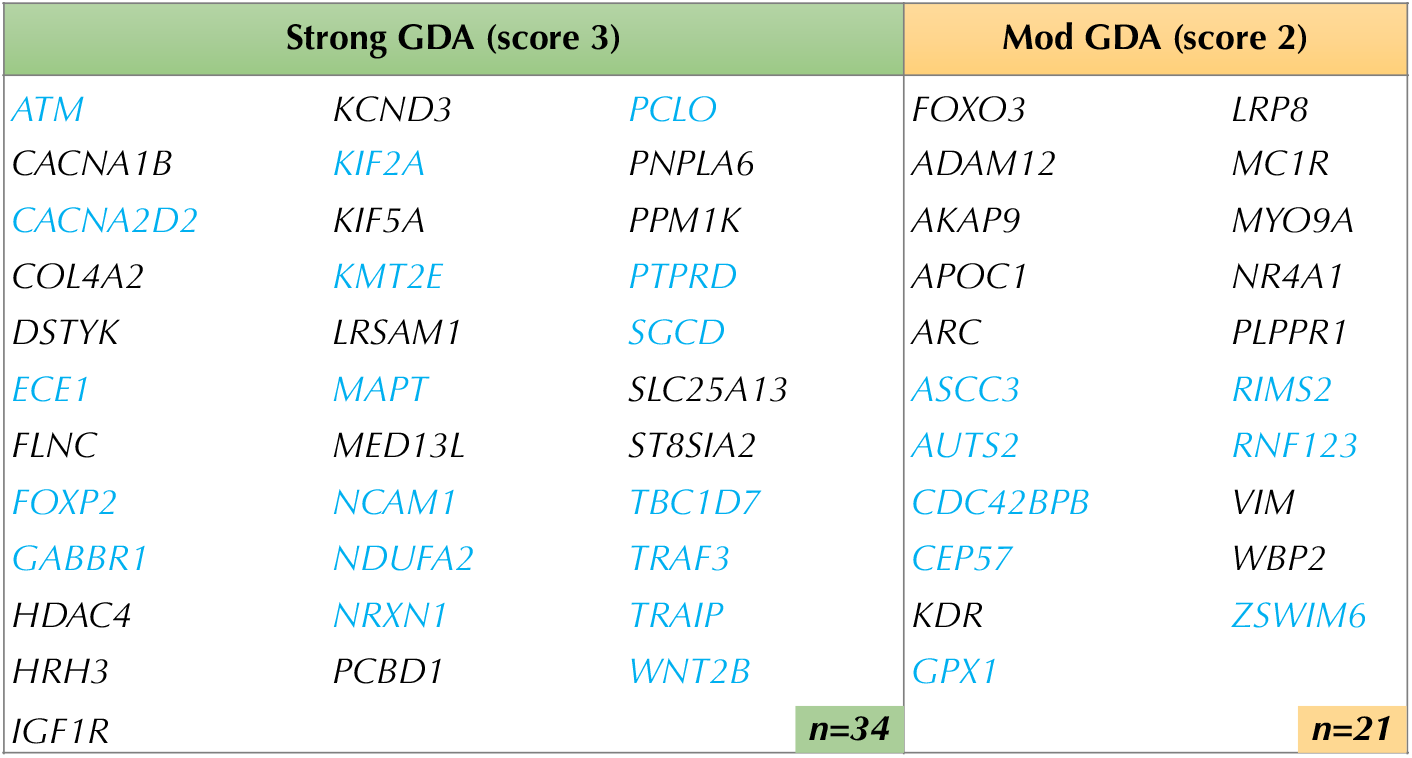
Phase 2 Advanced 55 Brain Targets with Evidence of CNS Disease Linkages. 55 of the Phase 1 targets were advanced through phase 2 on the basis of having at least one DisGeNET CNS-relevant disease association score with an Effect Size ≥ 0.3. Targets in blue (n=26) are supported by <u>both</u> transcriptomic and genomic evidence of association to PTSD. *Strong GDA* Effect Size *(≥ 0*.*4) —> score 3. Moderate (Mod) GDA* Effect Size *(0*.*3-0*.*39) —> score 2*.

**Fig. 3.**
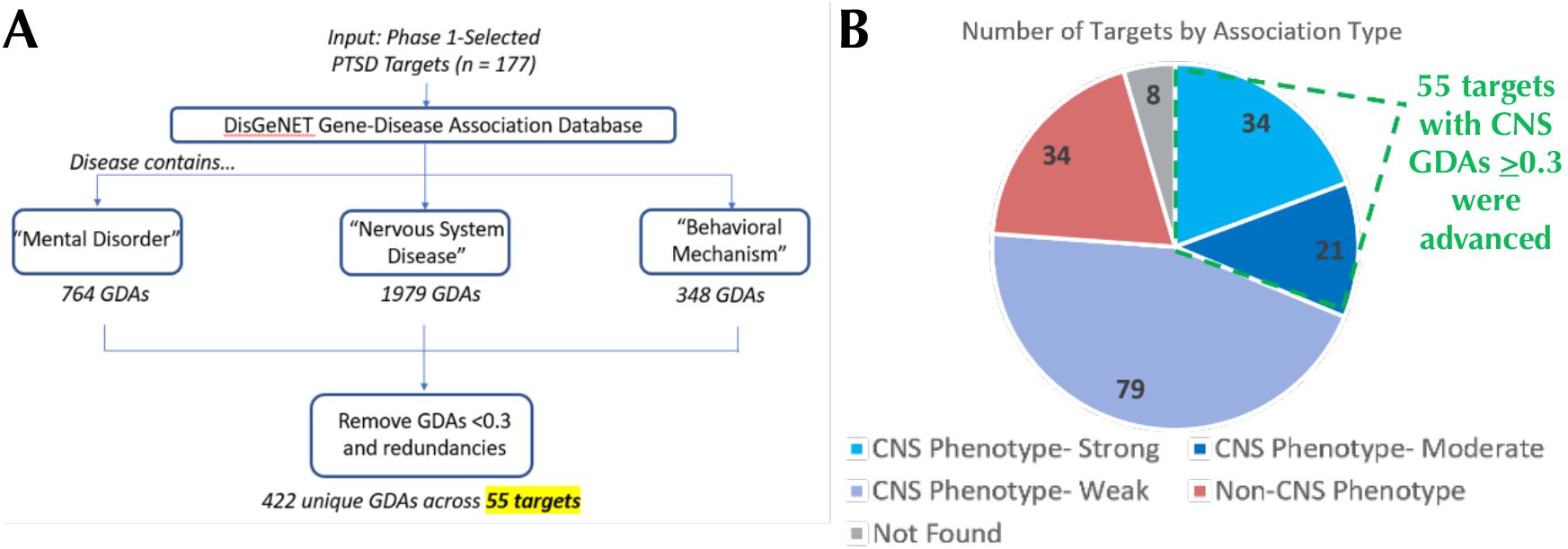
Phase 2 Leverages DisGeNET GDA Scores to Identify PTSD Targets with CNS Phenotype Linkages. For each Phase 1-advanced target, DisGeNET gene-disease association (GDA) scores for CNS-relevant disorders were exported and filtered to retain only GDA scores of ≥ 0.3 (**A**). 135 of the 177 phase 1-advanced targets had a CNS-relevant GDA score of any strength, while 55 targets had at least one moderate (0.3-0.39) or strong (≥ 0.4) CNS-relevant GDA score (**B**).

**Fig. 4.**
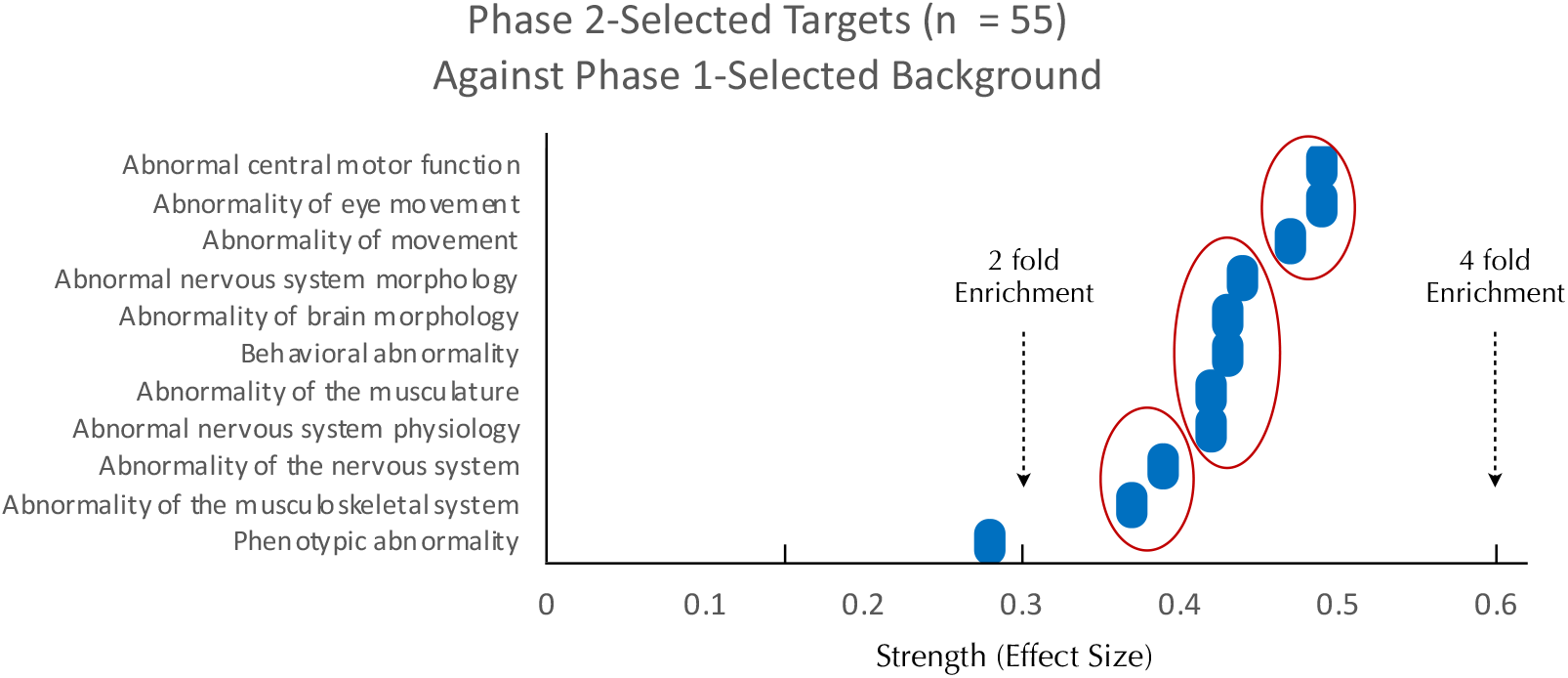
Phase 2-Selected Targets are Enriched for CNS Disease Above and Beyond Phase 1-Selected Targets. Panels represent all significantly enriched (FDR <0.05) terms resulting from Monarch human phenotype enrichment analysis via STRING. Phase 2 targets were enriched for CNS phenotypic abnormalities when tested against a stringent background of Phase 1-selected targets. “Strength” on the x-axis represents the effect size = log10[(Observed number or proteins in the analyzed network annotated to phenotype ‘X’:Total number of all proteins annotated to phenotype ‘X’):(Expected number of proteins annotated to phenotype ‘X’ in a random network of the same size)]. For perspective, a 2-fold enrichment would be equivalent to an effect size of 0.3.

This analysis supports that selecting targets in Phase 2, on the basis of CNS-relevant DisGeNET GDA scores, had the intended effect of enriching network for CNS-relevant pathology.

<u>The goal of phase 3</u> was to rank order targets with a composite pathogenicity score (**Fig. 1**), which was comprised of three components termed “Metrics” (**Fig. 5**), which are described below. Each Metric had a maximum score of 1, which yielded a maximum pathogenicity score of 3 that was used along with GDA scores to rank order PTSD targets (described below).

**Fig. 5.**
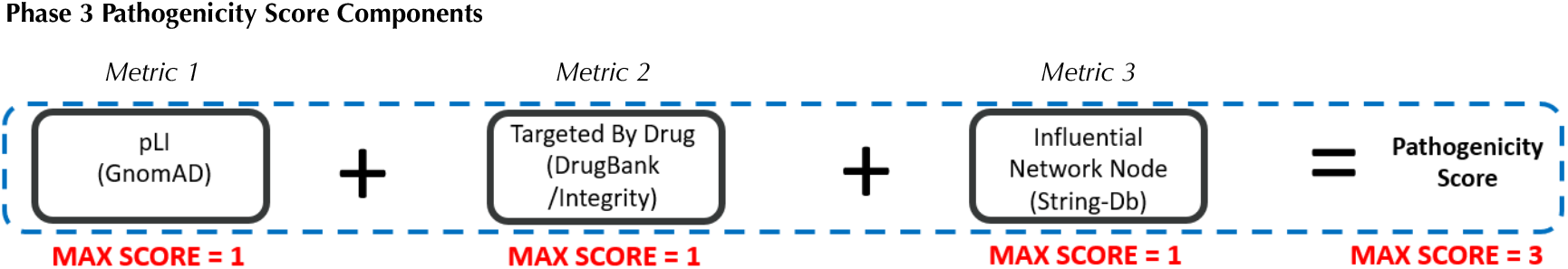
Phase 3 Pathogenicity Score Components. Phase 3 derived scores for equally weighted components (metrics) to calculate a pathogenicity score for each target. *Metric 1*: Predicted loss-of-function intolerance (pLI), *Metric 2*: Targeted by drug, and *Metric 3*: Influential network node.

### [Metric 1] Predicted loss-of-function intolerance (pLI)

<u>P</u>redicted loss-of-function intolerance (pLI) is a continuous metric with a range of 0 to 1 that quantifies the selection pressure against (i.e., the relative rarity of) loss-of-function (LOF) variants for a particular gene^**37**^.

A pLI score >0.9 is considered to be extremely loss-of-function intolerant and likely to be haploinsufficient; that is when one functional copy of a gene is not capable of rescuing a phenotype^**37**^. Importantly, the reason certain genes are loss-of-function intolerant is that these genes tend to be essential for normal cellular function, such that LOF results in severe, if not fatal phenotypes^**37**^. Therefore, pLI provides strong *positive* evidence that a putative target is pathogenic. However, it is important to stress that low a pLI score does not effectively rule out a target as benign^**37**^. For each Phase 2-advanced target pLI, reported by GnomAD/DisGeNET was used with no additional transformation (**Fig. 6**).

**Fig. 6.**
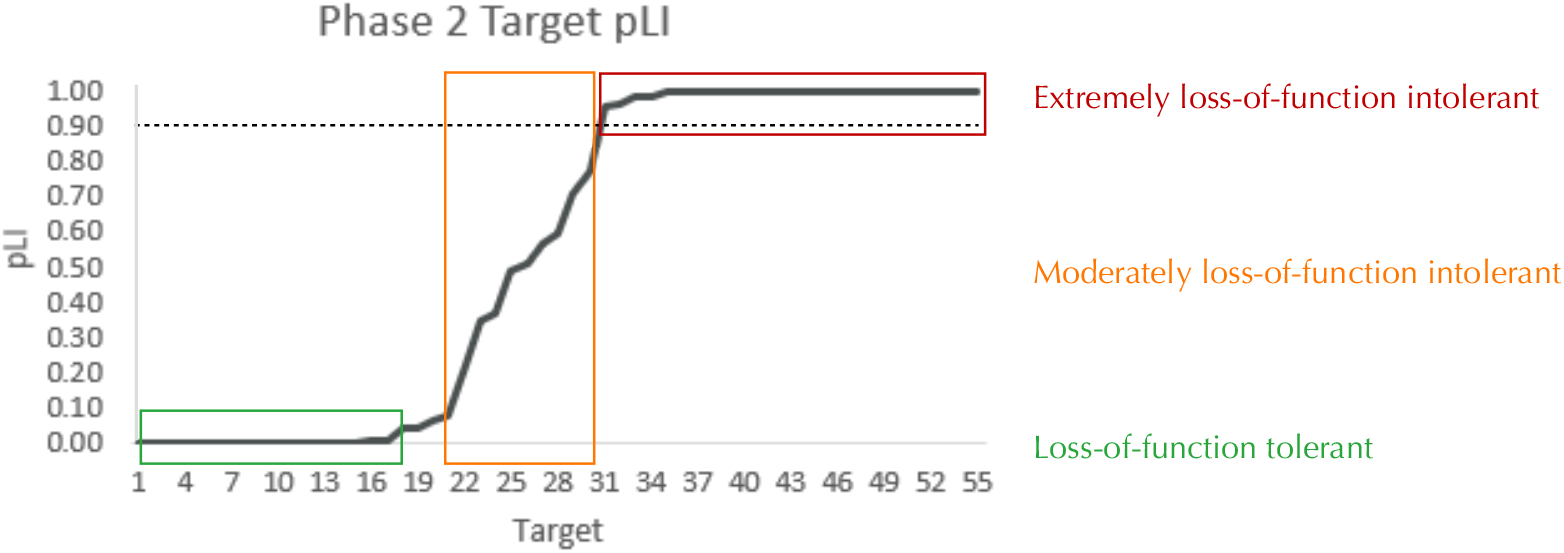
pLI among Phase 2-Advanced Targets. The predicted loss-of-function intolerance (pLI: 0-1) for each of the 55 phase 2-advanced targets. See **Supplemental Table 5** for detailed information on target-specific pLI. The dashed line indicates the standard threshold for a gene to be considered extremely loss-of-function intolerant (0.9).

### [Metric 2] Targeted by Drug

Drug information can inform estimates of a given target’s pathogenicity by providing *positive* evidence that modifying the drug target alleviates (portions of) the disease state. To generate a “targeted by drug” score for each target, drug information was mined from two sources: DrugBank (www.drugbank.com) and a legacy version of the Integrity Discovery Database. To estimate confidence that manipulating a target could be therapeutically effective, each target was scored on whether it has been targeted by a drug with a CNS indication and how advanced in development the drug was (**Table 2**). Among the 55 targets that we advanced from Phase 2, twelve targets had therapeutic drugs developed for CNS indications; the full scoring of these 12 targets is shown in **Table 3**.

**Table 2.** “Targeted by Drug” Scoring Schema. Based on current drug information, each target was evaluated to determine if a drug that modifies it is in development for a CNS indication and how far advanced in the development process it is.

|  | Score 0 | Score 0.25 | Score 0.375 | Score 0.5 |
| --- | --- | --- | --- | --- |
| CNS indication | No | n/a | n/a | Yes |
| Best CNS Phase | Preclinical | Phase 1 or 2 | Phase 3 | Launched |

### [Metric 3] Influential network node

Network analysis considers physical and biochemical interactions among selected targets and can help identify targets that reside in an ‘influential’ position by quantifying target connectivity within a disease-relevant network. To establish network nodes, the STRING database analysis tool^**21**^ was used to first generate a PTSD-relevant interaction network using the 177 targets advanced from Phase 1 (**Fig. 7**). Second, each of the 55 targets advanced from Phase 2 (represented as green nodes in **Fig. 7**) were annotated based on connectivity, where node degree is the number of proteins in the network with which a target interacts within the PTSD interaction network. The network node scores were assigned as a function of node degree with 0 degrees = 0, 1-3 degrees = 0.5, 4-5 degrees = 0.75, and >5 degrees = 1 (**Fig. 8**).

**Fig. 7.**
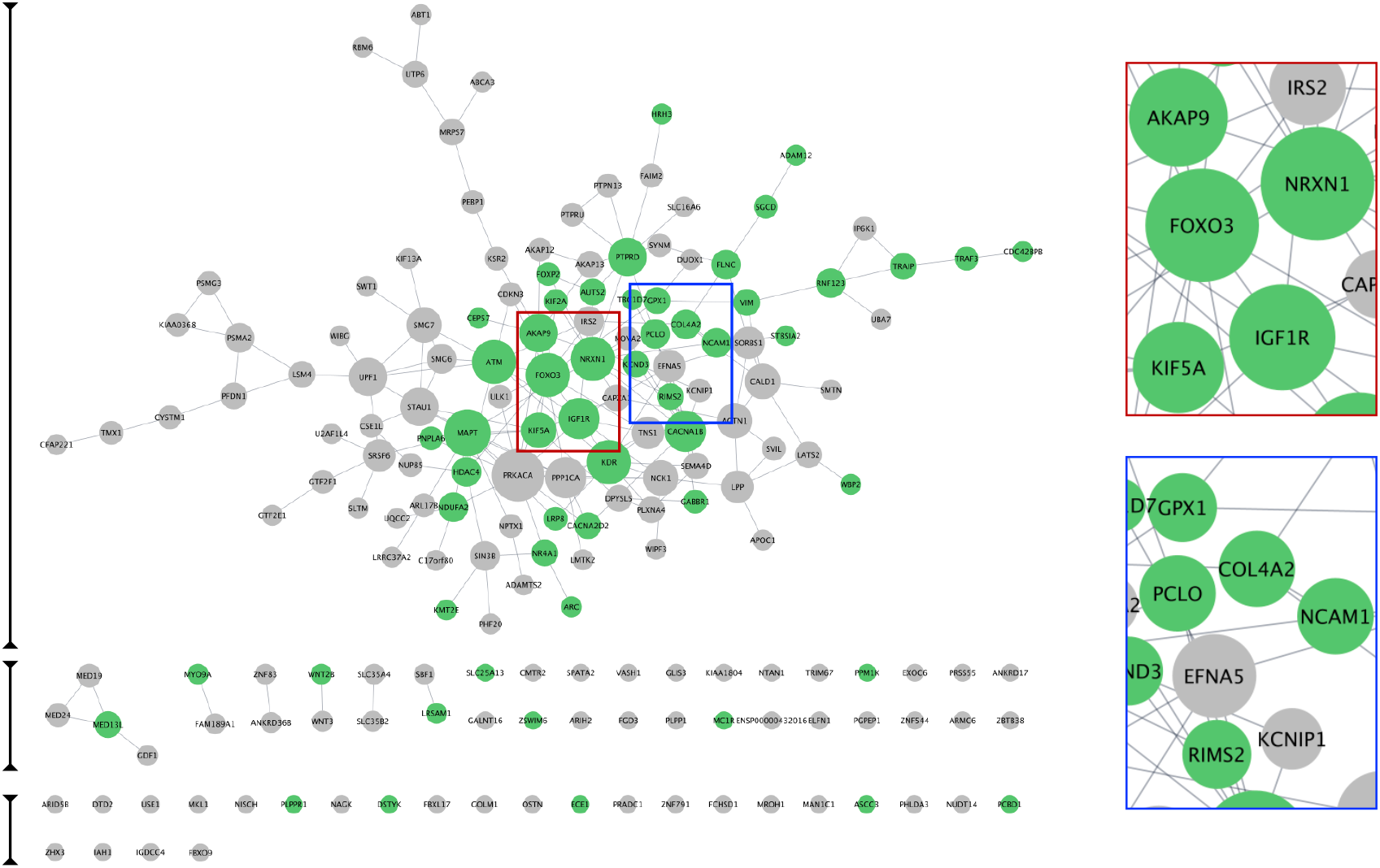
PTSD Target Interaction Network. STRING was used to generate a PTSD interaction network based on the 177 high confidence PTSD targets advanced from phase 1. The subset of 55 targets advanced by phase 2 are indicated as green nodes. Node size reflects the node degree, such that larger node size indicates greater node degree and hence more connections with other nodes. Two examples show examples of larger (red marque) and smaller (blue marque) nodes.

**Fig. 8.**
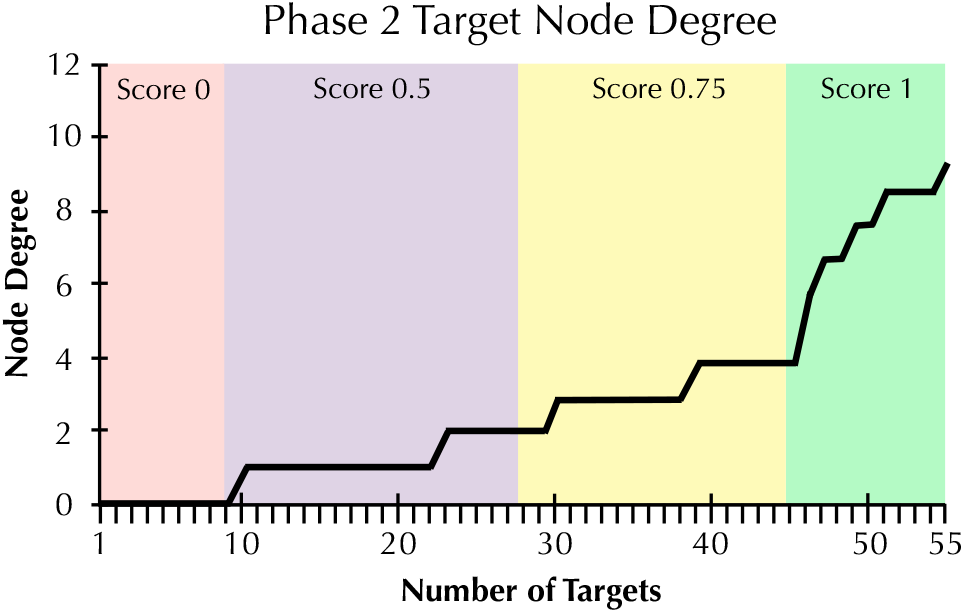
Node Degree among Phase 2-Advanced Targets. The node degree for each of the 55 phase 2-advanced targets within the 177-target phase 1 network. Scoring boundaries corresponding to target node degree are labeled. See **Supplemental Table 5** for details. Node Degree is the number of proteins in the network with which a target interacts.

### Integrating Phase 2 and Phase 3: Prioritizing Top Tier and Secondary (Ancillary) PTSD Targets

To generate a rank-ordered target priority list, target scores from Phase 2 (GDA Score) and Phase 3 (Pathogenicity Score) were summed to yield a novel <u>T</u>arget <u>A</u>dvancement <u>P</u>rioritization (TAP) Score (**Table 4**). Both the directionality of target changes in PTSD and the TAP score are valuable considerations for designing experiments that encompass our proposed Decision Matrix (**Fig. S1**). The top transcriptomics-based targets we identified include receptors (e.g. *GABBR1, HRH3, IGF1R*), ion channels (e.g. *CACNA1B, CACNA2D, KCND3*), epigenetic regulators (e.g. *HDAC4, KMT2E*), transcriptional regulators (e.g. *MED13L, FOXP2, ARC*), neurite regulators (*NRXN1, PCLO, MAPT, NCAM1*), and mediators of proteolysis (e.g. *ECE1, TRAF3*). Notably, 5 of the top 10 targets were supported by *both* transcriptomic and GWAS-derived evidence (**Supplemental Table 5**).

**Table 3.** “Targeted by Drug” Scores of Phase 2-Advanced Targets. The 12 (out of 55) targets that had at least one targeting compound in-development for a CNS indication were scored based on the criteria described in Table 3. The remaining 43 targets received a score of 0.

| Table 3<br>Targeted by Drug Score |  |  |  |
| --- | --- | --- | --- |
| Target | CNS Ind | CNS Phase | Score |
| *CACNA1B* | 0.5 | 0.5 | 1 |
| *CACNA2D* | 0.5 | 0.5 | 1 |
| *ECE1* | 0.5 | 0 | 0.5 |
| *GABBR1* | 0.5 | 0.5 | 1 |
| *GPX1* | 0.5 | 0.5 | 1 |
| *HDAC4* | 0.5 | 0.5 | 1 |
| *HRH3* | 0.5 | 0.5 | 1 |
| *IGF1R* | 0.5 | 0.375 | 0.875 |
| *KCND3* | 0.5 | 0.5 | 1 |
| *KDR* | 0.5 | 0 | 0.5 |
| *MC1R* | 0.5 | 0.5 | 1 |
| *NDUFA2* | 0.5 | 0.375 | 0.875 |

**Table 4.** Target Advancement Prioritization (TAP). From 177 targets, the top 20 transcript-based targets were rank ordered based on TAP score and partitioned into primary (n=10) and ancillary (n=10) targets, which serve as nodes. We also aligned targets based on CNS cell types and expression levels (transcripts per million) using GTeX. Directionality of target regulation in PTSD versus controls are shown with colored arrows. General function is also indicated. <u>Note</u>: NDUFA2 was swapped for FOXP2 due to low expression in neurons. Biological rationale for each target is briefly summarized.

| Target | Expression (transcripts per million) |  |  |  | Regulation and TAP |  | Biological Rationale in PTSD |
| --- | --- | --- | --- | --- | --- | --- | --- |
|  | Excitatory | Inhibitory | Astrocytes | Microglia | Direction (↑↓) | TAP | General Function |
| CACNA1B | 375.5 | 316.8 | 41.2 | 17.0 | ↑ | 5.999 | Ca<sup>2+</sup> ion channel, CNS disorders |
| IGF1R | 231.4 | 186.7 | 49.6 | 234.4 | ↑ | 5.842 | Receptor (TRK), CNS disorders |
| HDAC4 | 114.0 | 81.4 | 58.8 | 36.7 | ↑ | 5.750 | Deacetylase, CNS disorders |
| CACNA2D | 7.5 | 34.6 | 4.0 | 1.8 | ↑ | 5.749 | Ca<sup>2+</sup> ion channel, CNS disorders |
| KCND3 | 102.1 | 84.6 | 17.3 | 0.9 | ↑ | 5.739 | Kv<sup>+</sup> ion channel, CNS disorders |
| GABBR1 | 56.7 | 65.2 | 142.7 | 18.8 | ↑ | 5.499 | GABA<sub>B</sub> receptor, CNS disorders |
| NDUFA2 | 6.3 | 7.3 | 9.9 | 5.4 | ↓ | 5.398 | Dehydrogenase, NADH complex |
| PTPRD | 4422 | 1105 | 188.9 | 81.4 | ↑ | 5.000 | Phosphatase, CNS disorders |
| KIF5A | 27.5 | 20.8 | 2.8 | 0.0 | ↑ | 4.999 | Kinesin, CNS disorders |
| NRXN1 | 3095 | 2016 | 3818 | 166 | ↑ | 4.999 | Transmembrane R, CNS disorders |
| MED13L | 301.0 | 307.4 | 288 | 408 | ↑ | 4.750 | Scaffold, WNT,SHH, CNS disorders |
| PCLO | 632.9 | 715.2 | 38.4 | 48.3 | ↑ | 4.750 | Presynaptic scaffold, CNS disorders |
| NCAM1 | 375.5 | 316.8 | 503.5 | 34.0 | ↑ | 4.749 | Cell adhesion, CNS disorders |
| FLNC | 0.2 | 1.0 | 0.6 | 0.0 | ↑ | 4.749 | Attachment protein, CNS disorders |
| HRH3 | 4.1 | 22.7 | 0.0 | 0.0 | ↑ | 4.541 | Histamine R, memory impairment |
| KMT2E | 154.8 | 142.5 | 102.8 | 142.2 | ↑ | 4.500 | Methyl transferase, CNS disorders |
| FOXP2 | 179.9 | 6.1 | 5.3 | 275.5 | ↑ | 4.499 | Transcription factor, CNS disorders |
| KIF2A | 54.7 | 61.8 | 51.7 | 25.0 | ↑ | 4.499 | Kinesin, CNS disorders |
| KDR | 0.1 | 1.3 | 0.0 | 7.2 | ↑ | 4.499 | VEGF R (TRK), CNS disorders |
| TRAF3 | 104.7 | 60.5 | 51.7 | 22.4 | ↑ | 4.498 | Ubiquitin ligase, CNS disorders |

Targets indicated in blue (n=11) are supported by both transcriptomic & genomic evidence of association to PTSD
Primary Targets (n=10)
Ancillary Targets (n=10)

### PTSD target interaction network

As a next step we explored the interactions of our top 20 PTSD index proteins, which serve as nodes in our network. Nodes, which are connected by ‘edges’ reveal relations to each other and were used to essentially build the PTSD PPI network. To accomplish this, all 177 Phase 1 proteins were entered into STRING using the multiple proteins search tool and interrogated using STRING functions including the <u>M</u>arkov <u>Cl</u>uster (MCL) algorithm (**Fig. S2**), which revealed unique clusters. Subsequently, we imported our proteins into Cytoscape 3.10.2 to visualize, interrogate, and expand the PTSD and KCND3), all of which were on our Top 10 primary list and are involved in brain function. The second ranked target, which had a TAP score = 5.842 was IGF1R, which is a receptor tyrosine <u>k</u>inase (RTK) that is also involved in CNS disorders (**Table 4**). IGF1R had eight ‘first neighbors’ all of which were on our Phase-1 Advanced Target list (**Supplemental Table 4**), but only one appeared on our refined list of Top 20 targets (**Table 1**). Some of IGF1R first neighbor proteins were involved in brain function (e.g. axon guidance) as well as other functions including regulating phosphorylation and RAS signaling (**Fig. 10**). A third target (ranked 10) from our top 10 primary target list was NRXN1, which is a transmembrane receptor involved in CNS disorders (**Table 4**). First neighbor analysis revealed eight first neighbor proteins (NCAM1, RIMS2, FBXL17, FOXP2, PTPRD, AUTS2, CACNA1B, and MAPT), all of which have brain-specific functions (**Fig. 11A-C**). We also assessed NRXN1 edges in detail, which showed that edges (connections between nodes) were supported by both experimental and co-expression interactions (**Fig. 11D**). Cytoscape also revealed that NRXN1 interacts with numerous synaptic cleft proteins (**Fig. 11E**) (Rudenko, 2019).

**Fig. 9.**
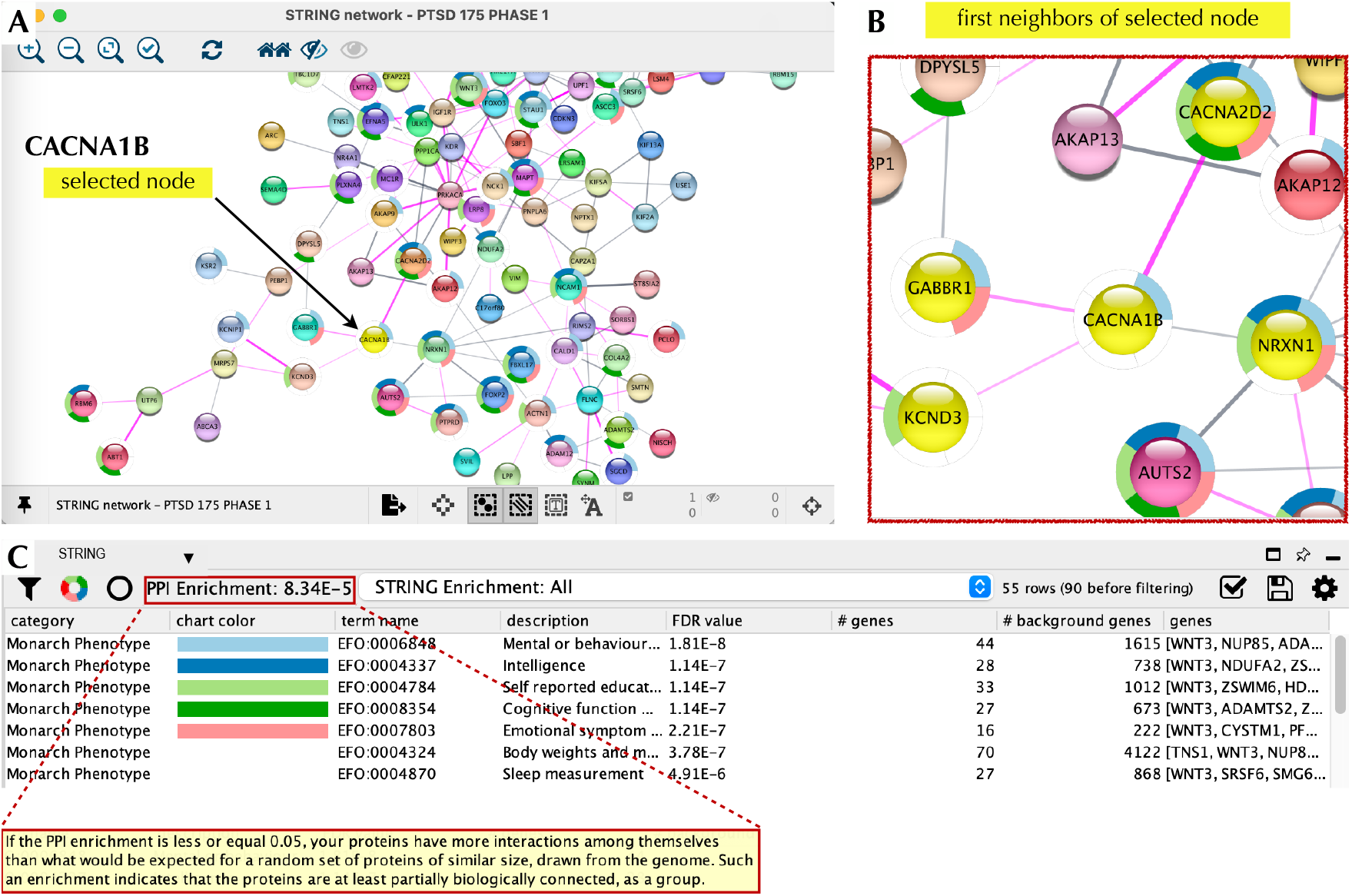
CACNA1B Node and Local Interaction Network. Cytoscape 3.10.2 was used to interrogate the PTSD interaction network based on CACNA1B as the node of interest (**A**). We then used the ‘first neighbors of selected node’ tool to generate a local interaction network, which revealed the presence of multiple interacting proteins, which were also part of our top 20 list (**B**). The ‘STRING enrichment’ tool showed that these proteins had a PPI enrichment of 8×10^−5^ and a role in brain health/function with a FDR <10^−7^ (**C**).

**Fig. 10.**
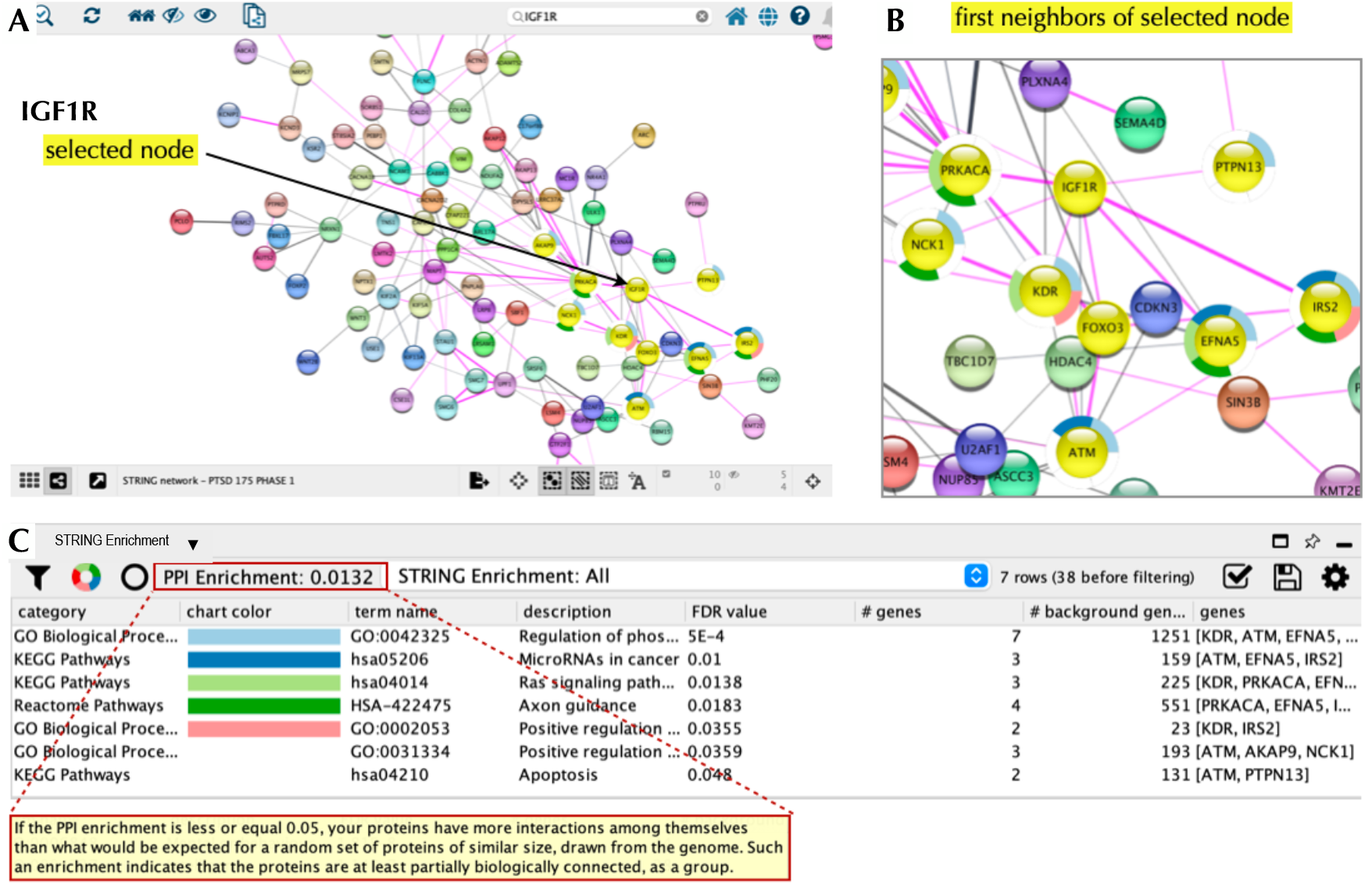
IGF1R Node and Local Interaction Network. Cytoscape 3.10.2 was used to interrogate the PTSD interaction network based on IGF1R as the node of interest (**A**). We then used the ‘first neighbors of selected node’ tool to generate a local interaction network revealing eight interacting proteins, which were also part of our top 20 list (**B**). The ‘STRING enrichment’ tool showed that these proteins had PPI enrichment and a role in stem cell differentiation, axonal path finding, and dendritic outgrowth^**55**^ with a FDR <0.05 (**C**).

**Fig. 11.**
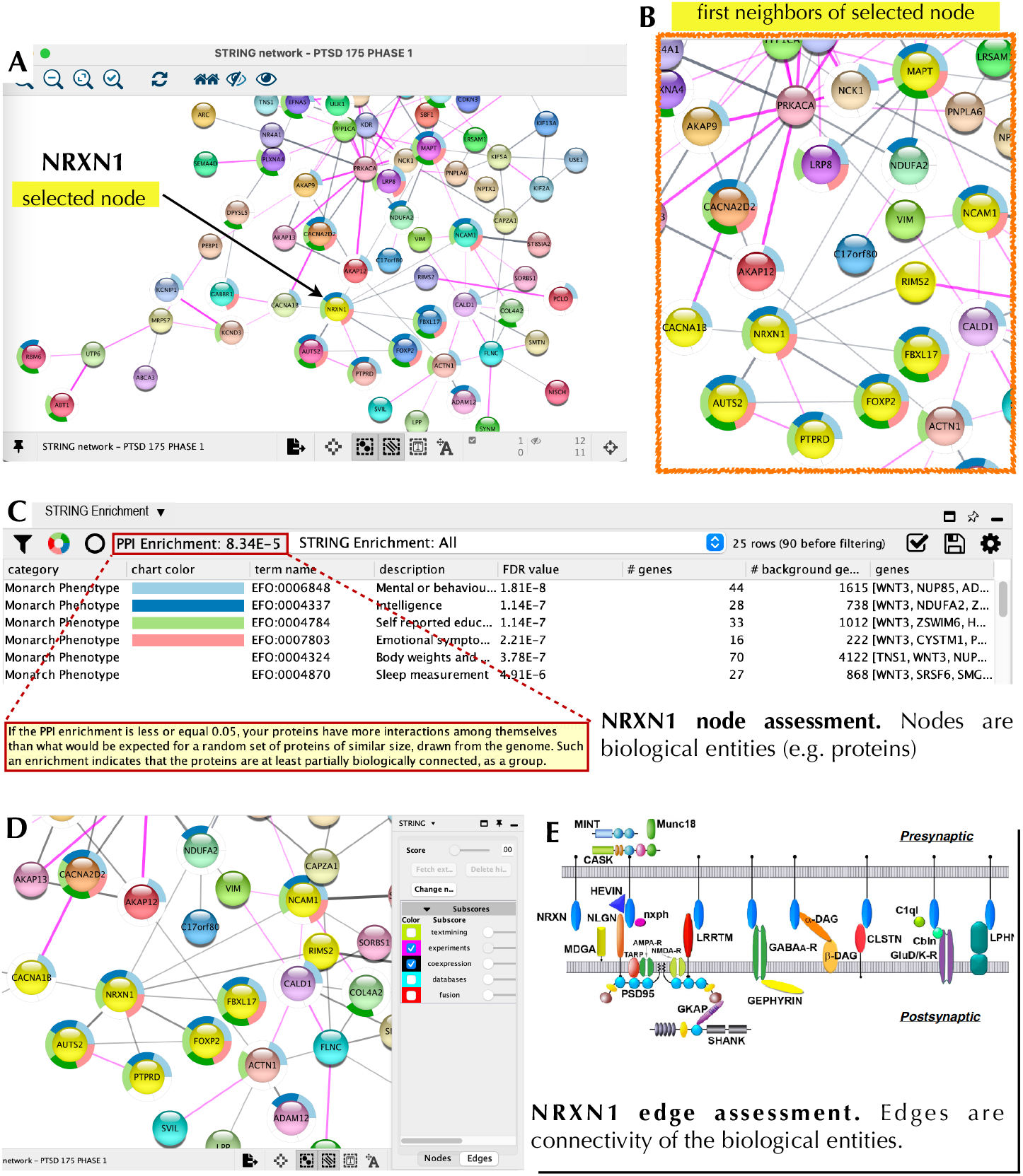
NRXN1 Node and Local Interaction Network. Cytoscape 3.10.2 was used to interrogate the PTSD interaction network based on NRXN1 as the node of interest (**A**). We then used the ‘first neighbors of selected node’ tool to generate a local interaction network revealing eight interacting proteins, which were also part of our top 20 list (**B**). The ‘STRING enrichment’ tool showed that these proteins had a PPI enrichment of 8×10^−5^ and a role in brain health/function with a FDR <10^−7^ (**C**). The panels **A-C** show node assessment while panel **D** shows edge assessment with subscores relevant to supporting evidence based on experiments & co-expression. The panel in the bottom right (**E**) shows NRXN1 (blue ovals) interacting with numerous distinct partners in the synaptic cleft (Rudenko, 2019).

PPI network using the Cytoscape ‘First Neighbor’ tool (**Fig. 9**). Our top target based on a TAP score of 5.999 (out of 6) was CACNA1B, which is a Ca2^+^ ion channel involved in CNS disorders (**Table 4**). CACNA1B had four ‘first neighbors’ (CACNA2D2, NRXN1, GABR1,

## DISCUSSION

PTSD is a significant and complex disorder with few therapeutic options, which notably have not been updated in over twenty years. GWAS have been employed to identify genetic targets of disease and substantial efforts and resources have been invested in PTSD GWAS, which have revealed that there is a heritable, and thus genetic contribution to the disorder^**5–8**^. However, it noteworthy that GWAS have typically not led to novel therapeutic targets nor a deeper understanding of neurological disorders^**40**^. Although progress has been made to overcome GWAS limitations^**41–43**^, aligning GWAS SNPs to specific actionable genes still has significant challenges. Therefore, we developed a streamlined approach that relied on transcripts derived from published RNAseq studies^**13–16**^ applied to postmortem CNS tissue and iNeu^Exc^, from both PTSD and control samples allowing us to identify novel targets. We subsequently utilized publicly available tools to prioritize the transcriptome-based targets with relevance to PTSD.

We clustered 20 top tier targets based on Cytoscape 3.10.2 and placed them in context with each other (**Fig. 12**). Nodes with significant edges, that is connections, with other top 20 targets from our initial list (**Table 1**) are shown as lines connecting with an oval. Targets that had support from both transcriptomics and GWAS are indicated with blue font. Only FOXP2 (bold blue font) was observed in a previous internal (CVB) PTSD SNP-AG assessment and in the current transcriptomic-based identification of PTSD targets. Interestingly, four super (functional)-clusters (**Fig. 12, I-IV**) bind targets into a functional network although each putative target has a distinct function (**Fig. 12 and Table 4**). Cluster III contains three nodes (NCAM1, MAPT, and KIFA5). Although, MAPT was not derived from transcriptomics directly it appeared as a node via edges connected to other nodes in our analysis with Cytoscape. MAPT also appeared in a previous internal SNP-AG identification campaign. Cluster I was connected to Cluster II via connection between the CACNA1B and NRXN1 nodes.

**Fig. 12.**
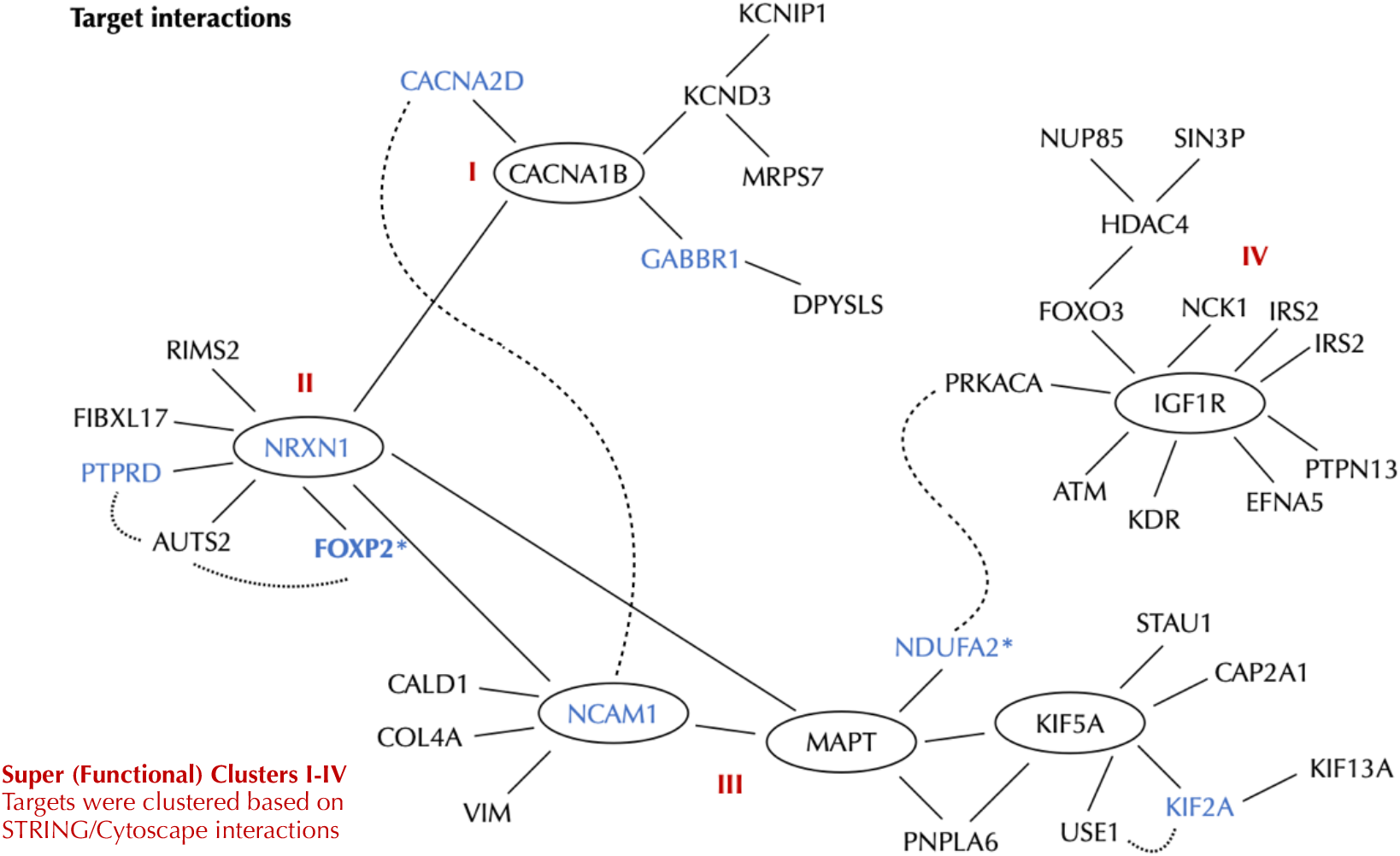
PTSD Targets and Local Interaction Network. Cytoscape 3.10.2 was used to interrogate the PTSD interaction network based on nodes, which were further organized into four super clusters (I-IV). The primary 10 ten targets are: CACNA1B, IGF1R, HDAC4, CACNA2D, KCND3, GABBR1, NDUFA2*, PTPRD, KIF5A, NRXN. The ancillary 10 targets are: MED13L, PCLO, NCAM1, FLNC, HRH3, KMT2E, FOXP2*, KIF2A, KDR, TRAF3. FOXP2* is an ancillary target and was swapped in as a primary target due to NDUFA2 having low expression in neurons (see Table 4). Targets with both transcriptomics & GWAS support are shown by blue font.

We denote Cluster II as a central hub with multiple edges (connections) that bind two nodes within Cluster III. Finally, Cluster IV initially appeared to be independent, but was revealed to be indirectly bound to Cluster III by edges that were connected by NDUFA2 and PRKACA. Notably, PRKACA was not identified via our top 20 list and thus represents an expansion component ascertained using Cytoscape and bridges the two clusters. Additionally the interactions of PTSD transcript nodes can be partitioned into four distinct Super (*functional*) clusters, which allow for more readily assessing both function and putative underlying pathobiology of PTSD. Nodes within functional clusters, which represent possible druggable targets, and their associated pathways, may be utilized to ameliorate cellular disease phenotypes.

<u>CACNA1B</u>, which is the node of Cluster 1, is closely associated with CACNA2D. Both are voltage gated calcium channels involved in neurotransmitter release and implicated in a number of CNS disorders including seizures, ataxia, and schizophrenia^**45**^. CACNA1B is connected to <u>NRXN1</u>, which is the node of Cluster II and is a cell surface receptor. Cell surface receptors represent traditional pharmacologically-druggable targets. NRXN1 regulates Ca^2+^ dependent processes including neurotransmitter release and synaptic function^**47,50**^. NRXN1 is connected to <u>NCAM1</u>, which is a cell adhesion molecule involved in cell-cell interactions and cell-matrix interactions. NRXN1 is also implicated in schizophrenia and bipolar disorder. <u>KIF5A</u>, part of Complex III, is a kinesin enzyme involved in axonal transport, as well as other functions, and is implicated in neurological disorders^**56**^. <u>IGF1R</u>, the node within Cluster IV is a receptor tyrosine kinase, which is another traditional pharmacologically-druggable target involved in cell growth and survival. IGF1R is implicated in both Alzheimer’s disease and Parkinson’s Disease^**55**^.

The ten primary targets identified using our approach (CACNA1B, IGF1R, HDAC4, CACNA2D, KCND3, GABBR1, NDUFA2*, PTPRD, KIF5A, NRXN) were expanded and clustered revealing broadly related functional interactions that at face-value help distinguish underlying biological processes disrupted as a result of PTSD. In addition to the convergence of functionality there are neurological diseases related to the clusters that have strong affiliation with PTSD and at a cellular level suggests overlap between mechanisms contributing to PTSD and other neurological disorders. Interestingly, NDUFA2 is expressed at very low levels in neurons and was the rationale for swapping FOXP2 from our ancillary target list in place of NDUFA2. Additionally, FOXP2 is expressed highly in excitatory neurons and is also implicated in schizophrenia, autism, and major depressive disorder.

Our primary goal was to identify novel PTSD targets and concomitantly uncover mechanisms that contribute to the disorder. However, our target prioritization strategy may also be applied to a wide array of diseases to accommodate the need for additional or new therapeutic drug targets and to facilitate the procurement of cellular disease phenotypes prior to initiating drug discovery efforts. Of course, other cell types and phenotypic classes of interest (e.g., immune or cardiovascular disorders) may be emphasized through choice of input datasets and disease class filtering in DisGeNET. Importantly, as cell-type specific datasets and datasets from additional brain regions become more available, it is anticipated that cell/tissue type of observations will become a more significant variable for Phase I stage prioritization. Other types of biological data may also be incorporated as supporting evidence, such as DNA methylation.

In addition, genotype-phenotype linkage information generated as part of a prioritization process can be mined for hypothesis generation and functional interpretation. For example, recently published papers describe several related approaches that leverage gene-disease associations maps similar to what was generated in our Phase 2 strategy (described above) to make mechanistic predictions and stratify complex diseases^**26,38,39**^.

The rationalized and systematic prioritization strategy described and applied here nominated 46 (i) Independently replicated PTSD-associated targets with(ii)Observed differential expression in PTSD brain tissue/excitatory neurons and (iii) Evidence supporting CNS-relevant pathogenicity. The PTSD targets we prioritized may be advanced to *in vitro* experimental functional validation by following our rationalized Decision Matrix (**Fig. S1**), which is anticipated to improve confidence in observing phenotypic effects in specific subsets of neurons.

There are alternative methods describing the establishment of PPI networks^**57**^, which may result in more robust networks, but the tools we utilized are free and easy to use and allowed us to establish a PTSD network for functional interrogation. We specifically undertook this task because for PTSD there are no new therapies with the existing class of SSRI therapeutics being developed two decades ago. This is particularly relevant given that a promising ‘psychedelic’ molecule (MDMA) recently did not receive approval for PTSD^**58**^. This suggests a need to continue to identify novel targets to both understand this complex disorder and develop therapeutic alternatives to complement or supplement SSRIs or psychedelics, provided the former are approved for treating PTSD. Here we provide roles of the associated networks in contributing to PTSD.

### Functional Cluster I

CACNA1B (L-type) and CACNA2D (non-L-type) (**Fig. 12**) belong to voltage gated calcium channels, which modulate diverse functions in the CNS. CACNA1B and GABBR1 are enriched in the GABAergic <u>synapse</u> pathway and are up-regulated following propofol (anesthetic) exposures^**44**^. CACNA1B and GABBR1 are also involved in biological pathways and mechanisms associated with genes implicated in schizophrenia^**45**^. CACNA2D is a risk gene for another complex disorder, Autism^**46**^. Because both calcium channel types converge on phenotypic associations with PTSD (**Table S4**), this provides reinforcing evidence that these channels contribute broadly to complex neurological disorders.

### Functional Cluster II

It has been suggested that FOXP2, DISC1 and the NRXN family are linked in a molecular network that contributes to neurodevelopmental disorders^**47**^. However, this is the first report that we are aware of that provides supporting evidence that NRXN1 and FOXP2 are interacting partners (**Fig. 12**). It appears that FOXP2 and PTPRD are both involved in maintaining <u>synaptic</u> architecture, which emerged only after complex mutagenesis assessment^**48,49**^. In addition, both proteins appear to form both pre-synaptic and post-synaptic structural complexes^**50**^.

### Functional Cluster III

NDUFA2 has previously been shown to be related to the Avoidance subdomain of the PTSD symptom cluster^**51**^. NDUFA2 is an element of the Mitochondrial complex 1 and is dysregulated in neurological disorders such as AD^**52**^. We could not find a direct link between MAPT and NDUFA2 (**Fig. 12**) in the literature so the interaction between them is considered tenuous at this time. Kinesins (also called KIFs) are molecular motors and implicated in both fast (50–400 mm/day) and slow (less than 8 mm/day) axonal transport^**53**^. KIF2A appears to be an essential regulator of neuronal connectivity and for the establishment of precise postnatal hippocampal wiring^**54**^. Thus, a plausible scenario is that perturbations in KIF2A and KIF5A result in altered interactions with TAU (encoded by *MAPT*) resulting in disrupted axonal transport as a contributing mechanism to PTSD.

### Functional Cluster IV

IGF-1 (**Fig. 12**) has a major role in neuronal development as it supports neuronal stem cell differentiation, axonal path finding, and dendritic outgrowth^**55**^. Supportingly, studies on the role of the IGF-1 receptor elicit very similar phenotype^**55**^. IGF-1 acts locally via IGF1R to augment <u>synaptic</u> connections based on olfactory nervous system assessment in mice^**55**^.

Collectively, the four nodes play a role in synaptic architecture and function as well as related axonal transport, which suggests that alterations in these processes contributes to PTSD and suggests cellular interactions to probe in discovery model systems. Finally, limitations to the study include the inability to partition putative targets based on distinct cell types including neuronal subtypes, astrocytes, and microglia. An additional limitation is the absence of non-cortical tissue representation including the amygdala and striatum.

#### A Path Ahead

Given the poor translational validity of preclinical models of PTSD (e.g. SPS^**62**^), which were evaluated through various programs and methods developed to improve their utility^**63**^ [EQIPD, GOT-IT, Emmerich, et al 2021] we propose, as a future direction, to validate these PTSD targets and interrogate their involvement in specific pathways using <u>human</u> iPSC models. While this section is hypothetical, we wanted to propose a logical strategic example for efficiently and cost-effectively validating targets (**Fig. S1**). Initially, proposed targets would be ascertained for their expression and distribution at the RNA/protein level using induced iPSC-derived <u>exc</u>itatory <u>neu</u>rons (iNeu^Exc^). Traditional iPSC-derived neurons, which are generated by taking cells with a somatic cell fate (e.g. fibroblasts) and reprogramming them to obtain an iPSC (stem cell) identity^**66,67**^. However, this process essentially eliminates disease phenotypes. Subsequently, stem cells are programed to obtain a unique somatic fate such as a subtype-specific neuron (e.g. cortical excitatory neuron). In contrast, a process of direct conversion from fibroblast to excitatory neuron (iNeu^Exc^) preserves disease phenotypes making them better suited to understand cellular disease pathology^**68**^. This is an important premise for using advanced iPSC technology to interrogate disease pathology occurring in adult neurons. The protocol for generating iNeu^Exc^ has been publicly available for years making them a suitable substrate to begin studies although it would also be valuable to follow the same process to produce induced iPSC-derived <u>Inh</u>ibitory <u>neu</u>rons (iNeu^Inh^).

Determination of expression and distribution are the first decision point with targets having validated expression being genetically manipulated in accordance to directionality (see **Table 4**). CRISPR modification is proposed to be coupled with subjecting iNeu^Exc^ to a stressor (**Fig. S1A**) such as oxytocin or hydrocortisone^**11,12**^. For PTSD, the goal is to mimic the convergence of genetic/transcriptomic perturbation plus an environmental stressor.

We complement our target identification and prioritization strategy with a Decision Matrix that describes the logic of functionally interrogating prioritized PTSD targets using relevant assays for PTSD target validation (**Fig. S1**). Morphometric and electrophysiological assays respectively, are informative or decision points with altered electrophysiology being viewed as a particularly important phenotype based our top tier target properties (**Table 4**). Tar gets demonstrating altered electrophysiological properties would be advanced through the Decision Matrix to gather valuable information to facilitate understanding of the biological alterations that underpin PTSD clinical phenotypes.

The next critical step would be to validate one or more of these targets. Validation of this methodology would provide Industry with a tool to make rational drug development decisions that would increase their likelihood of success, decrease unnecessary testing of patients in negative clinical trials, and speed the time to important therapeutic options. Among the 20 putative PTSD targets we identified, five have a combined ten compounds at various stages of clinical development (**Supplemental Table 8**). Notably, all ten compounds are being tested for, or are already approved for, one of two CNS indications, epilepsy and neurodegeneration. Two are approved for use in epilepsy and four other compounds of the same targets (CACN1B and HDAC) are also in development for epilepsy. Two other targets (GABBR1 and KDR) have one compound each being developed for epilepsy while the compounds targeting the fifth target, IGF1R, are being tested in neurodegeneration. An important next step in validating this methodology is to use one in the Decision Matrix (**Fig. S1**) to ameliorate electrophysiological phenotypes elicited by CRISPR modification. If applicable, this subset of targets could be considered for repurposing^**69**^ to treating PTSD.

In summary, we developed a novel strategy to identify and prioritize PTSD targets and place them in biologically meaningful pathways. In addition, we propose a strategic framework for validating biological properties. Our goal was to make these approached publicly available to benefit PTSD patients and researchers pursuing therapeutic development for the disorder, but we also propose that the tools can be utilized in other disease discovery contexts.

## ACKNOWLEDGMENTS

We acknowledge Elena Rotondo Engeldrum, Ph.D. for her significant contribution to developing the transcriptomics strategy and conducting essential analyses. MZ initiated and contributed to the transcriptomics strategy. ERE and MZ developed the local interaction network, wrote the manuscript, and generated figures. ERE, MZ, AG, MH provided equal intellectual contribution. The time and contribution of authors and acknowledged staff occurred exclusively while employees at Cohen Veterans Bioscience. ERE is currently at Regeneron.

## SUPPLEMENTAL MATERIAL

**Supplemental Table 1. PTSD_Brain_Target_Prioritization _Phase1**

**Supplemental Table 2. PTSD_Brain_Target_Prioritization_Phase2**

**Supplemental Table 3. PTSD_Brain_Target_Prioritization_Phase3**.

**Link for Tables 1-3:** https://drive.google.com/drive/folders/1FaG0E_5ConmyUvuuV_LznwsHQjoxpoJs?usp=sharing

**Supplemental Table 4.** Phase 1 Advanced 177 Brain Targets that are Confidently Associated to PTSD. 177 targets were prioritized on the basis of direct observation of differential expression in PTSD brain tissues or iPSC-neurons, replication in independent cohorts, and consistent direction of observed difference relative to controls. Targets in blue (n=65) are supported by both transcriptomic and genomic evidence.

| Supplemental Table 4<br>177 Phase-1 Advanced Targets (alphabetical order) |  |  |  |  |  |
| --- | --- | --- | --- | --- | --- |
| ABCA3 | CFAP221 | HDAC4 | MKL1 | PRADC1 | SWT1 |
| ABT1 | CMTR2 | HRH3 | MROH1 | PRKACA | SYNM |
| AC005697.1 | COL4A2 | IAH1 | MRPS7 | PRSS55 | TBC1D7 |
| AC244131.2 | CSE1L | IGDCC4 | MYO9A | PSMA2 | TMX1 |
| ACTN1 | CYSTM1 | IGF1R | NAGK | PSMG3 | TNS1 |
| ADAM12 | DPYSL5 | IP6K1 | NCAM1 | PTPN13 | TRAF3 |
| ADAMTS2 | DSTYK | IRS2 | NCK1 | PTPRD | TRAIP |
| AKAP12 | DTD2 | KCND3 | NDUFA2 | PTPRU | TRIM67 |
| AKAP13 | DUOX1 | KCNIP1 | NISCH | PYM1 | U2AF1L4 |
| AKAP9 | ECE1 | KDR | NOVA2 | RBM6 | UBA7 |
| ANKHD1-EIF4EBP3 | EFNA5 | KIAA0368 | NPTX1 | RIMS2 | ULK1 |
| ANKRD17 | ELFN1 | KIF13A | NR4A1 | RNF123 | UPF1 |
| ANKRD36B | EXOC6 | KIF2A | NRXN1 | SBF1 | UQCC2 |
| APOC1 | FAIM2 | KIF5A | NTAN1 | SEMA4D | USE1 |
| ARC | FAM189A1 | KMT2E | NUDT14 | SGCD | UTP6 |
| ARID5B | FBXL17 | KSR2 | NUP85 | SIN3B | VASH1 |
| ARIH2 | FBXO9 | LATS2 | OSTN | SLC16A6 | VIM |
| ARL17B | FCHSD1 | LMTK2 | PCBD1 | SLC25A13 | WBP2 |
| ARMC6 | FGD3 | LPP | PCLO | SLC35A4 | WIPF3 |
| ASCC3 | FLNC | LRP8 | PEBP1 | SLC35B2 | WNT2B |
| ATM | FOXO3 | LRRC37A2 | PFDN1 | SLTM | WNT3 |
| AUTS2 | FOXP2 | LRSAM1 | PGPEP1 | SMG6 | ZBTB38 |
| C17orf80 | GABBR1 | LSM4 | PHF20 | SMG7 | ZHX3 |
| CACNA1B | GALNT16 | MAN1C1 | PHLDA3 | SMTN | ZNF544 |
| CACNA2D2 | GDF1 | MAP3K21 | PLPP1 | SORBS1 | ZNF791 |
| CALD1 | GLIS3 | MAPT | PLPPR1 | SPATA2 | ZNF83 |
| CAPZA1 | GOLM1 | MC1R | PLXNA4 | SRSF6 | ZSWIM6 |
| CDC42BPB | GPX1 | MED13L | PNPLA6 | ST8SIA2 |  |
| CDKN3 | GTF2E1 | MED19 | PPM1K | STAU1 |  |
| CEP57 | GTF2F1 | MED24 | PPP1CA | SVIL |  |

**Supplemental Table 5.**
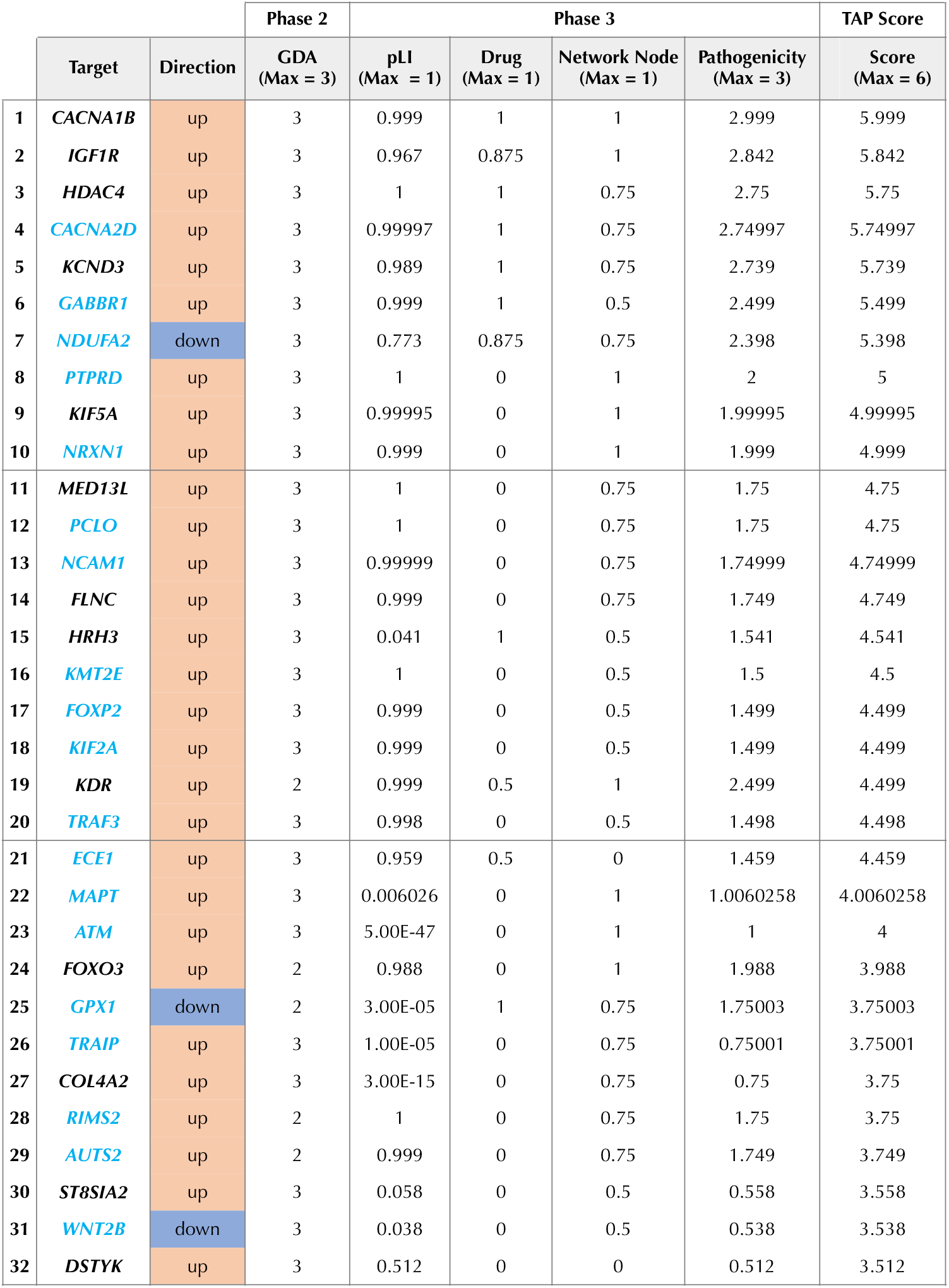
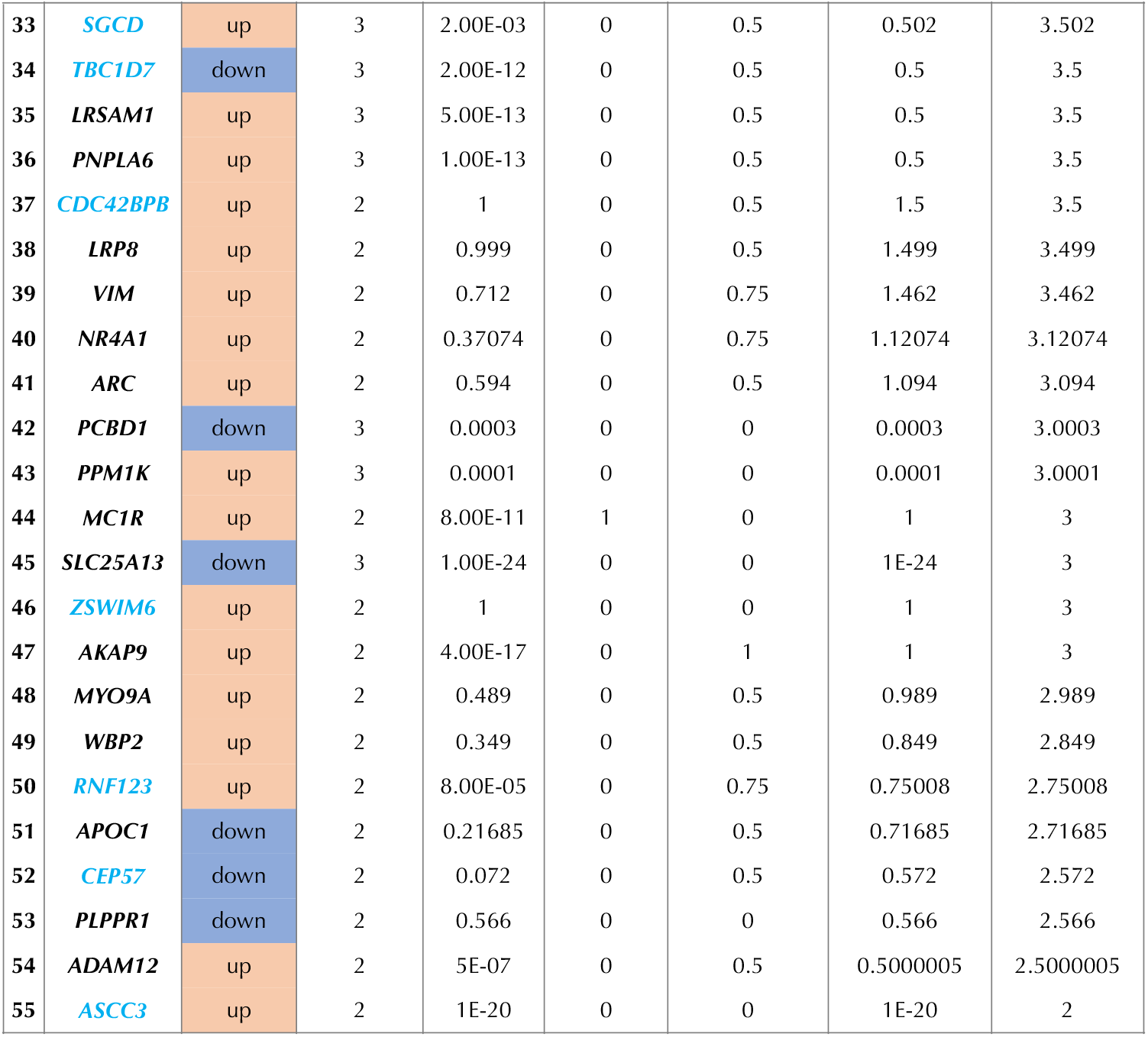
[<u>T</u>arget Advancement Prioritization (TAP) Score].

|  |  |  | Phase 2 | Phase 3 |  |  |  | TAP Score |
| --- | --- | --- | --- | --- | --- | --- | --- | --- |
|  | Target | Direction | GDA<br>(Max = 3) | pLI<br>(Max = 1) | Drug<br>(Max = 1) | Network Node<br>(Max = 1) | Pathogenicity<br>(Max = 3) | Score<br>(Max = 6) |
| 1 | *CACNA1B* | up | 3 | 0.999 | 1 | 1 | 2.999 | 5.999 |
| 2 | *IGF1R* | up | 3 | 0.967 | 0.875 | 1 | 2.842 | 5.842 |
| 3 | *HDAC4* | up | 3 | 1 | 1 | 0.75 | 2.75 | 5.75 |
| 4 | *CACNA2D* | up | 3 | 0.99997 | 1 | 0.75 | 2.74997 | 5.74997 |
| 5 | *KCND3* | up | 3 | 0.989 | 1 | 0.75 | 2.739 | 5.739 |
| 6 | *GABBR1* | up | 3 | 0.999 | 1 | 0.5 | 2.499 | 5.499 |
| 7 | *NDUFA2* | down | 3 | 0.773 | 0.875 | 0.75 | 2.398 | 5.398 |
| 8 | *PTPRD* | up | 3 | 1 | 0 | 1 | 2 | 5 |
| 9 | *KIF5A* | up | 3 | 0.99995 | 0 | 1 | 1.99995 | 4.99995 |
| 10 | *NRXN1* | up | 3 | 0.999 | 0 | 1 | 1.999 | 4.999 |
| 11 | *MED13L* | up | 3 | 1 | 0 | 0.75 | 1.75 | 4.75 |
| 12 | *PCLO* | up | 3 | 1 | 0 | 0.75 | 1.75 | 4.75 |
| 13 | *NCAM1* | up | 3 | 0.99999 | 0 | 0.75 | 1.74999 | 4.74999 |
| 14 | *FLNC* | up | 3 | 0.999 | 0 | 0.75 | 1.749 | 4.749 |
| 15 | *HRH3* | up | 3 | 0.041 | 1 | 0.5 | 1.541 | 4.541 |
| 16 | *KMT2E* | up | 3 | 1 | 0 | 0.5 | 1.5 | 4.5 |
| 17 | *FOXP2* | up | 3 | 0.999 | 0 | 0.5 | 1.499 | 4.499 |
| 18 | *KIF2A* | up | 3 | 0.999 | 0 | 0.5 | 1.499 | 4.499 |
| 19 | *KDR* | up | 2 | 0.999 | 0.5 | 1 | 2.499 | 4.499 |
| 20 | *TRAF3* | up | 3 | 0.998 | 0 | 0.5 | 1.498 | 4.498 |
| 21 | *ECE1* | up | 3 | 0.959 | 0.5 | 0 | 1.459 | 4.459 |
| 22 | *MAPT* | up | 3 | 0.006026 | 0 | 1 | 1.0060258 | 4.0060258 |
| 23 | *ATM* | up | 3 | 5.00E-47 | 0 | 1 | 1 | 4 |
| 24 | *FOXO3* | up | 2 | 0.988 | 0 | 1 | 1.988 | 3.988 |
| 25 | *GPX1* | down | 2 | 3.00E-05 | 1 | 0.75 | 1.75003 | 3.75003 |
| 26 | *TRAIP* | up | 3 | 1.00E-05 | 0 | 0.75 | 0.75001 | 3.75001 |
| 27 | *COL4A2* | up | 3 | 3.00E-15 | 0 | 0.75 | 0.75 | 3.75 |
| 28 | *RIMS2* | up | 2 | 1 | 0 | 0.75 | 1.75 | 3.75 |
| 29 | *AUTS2* | up | 2 | 0.999 | 0 | 0.75 | 1.749 | 3.749 |
| 30 | *ST8SIA2* | up | 3 | 0.058 | 0 | 0.5 | 0.558 | 3.558 |
| 31 | *WNT2B* | down | 3 | 0.038 | 0 | 0.5 | 0.538 | 3.538 |
| 32 | *DSTYK* | up | 3 | 0.512 | 0 | 0 | 0.512 | 3.512 |

|  | Target | Direction | Phase 2 | Phase 3 |  |  |  | TAP Score |
| --- | --- | --- | --- | --- | --- | --- | --- | --- |
|  |  |  | GDA<br>(Max = 3) | pLI<br>(Max = 1) | Drug<br>(Max = 1) | Network Node<br>(Max = 1) | Pathogenicity<br>(Max = 3) | Score<br>(Max = 6) |
| 33 | *SGCD* | up | 3 | 2.00E-03 | 0 | 0.5 | 0.502 | 3.502 |
| 34 | *TBC1D7* | down | 3 | 2.00E-12 | 0 | 0.5 | 0.5 | 3.5 |
| 35 | *LRSAM1* | up | 3 | 5.00E-13 | 0 | 0.5 | 0.5 | 3.5 |
| 36 | *PNPLA6* | up | 3 | 1.00E-13 | 0 | 0.5 | 0.5 | 3.5 |
| 37 | *CDC42BPB* | up | 2 | 1 | 0 | 0.5 | 1.5 | 3.5 |
| 38 | *LRP8* | up | 2 | 0.999 | 0 | 0.5 | 1.499 | 3.499 |
| 39 | *VIM* | up | 2 | 0.712 | 0 | 0.75 | 1.462 | 3.462 |
| 40 | *NR4A1* | up | 2 | 0.37074 | 0 | 0.75 | 1.12074 | 3.12074 |
| 41 | *ARC* | up | 2 | 0.594 | 0 | 0.5 | 1.094 | 3.094 |
| 42 | *PCBD1* | down | 3 | 0.0003 | 0 | 0 | 0.0003 | 3.0003 |
| 43 | *PPM1K* | up | 3 | 0.0001 | 0 | 0 | 0.0001 | 3.0001 |
| 44 | *MC1R* | up | 2 | 8.00E-11 | 1 | 0 | 1 | 3 |
| 45 | *SLC25A13* | down | 3 | 1.00E-24 | 0 | 0 | 1E-24 | 3 |
| 46 | *ZSWIM6* | up | 2 | 1 | 0 | 0 | 1 | 3 |
| 47 | *AKAP9* | up | 2 | 4.00E-17 | 0 | 1 | 1 | 3 |
| 48 | *MYO9A* | up | 2 | 0.489 | 0 | 0.5 | 0.989 | 2.989 |
| 49 | *WBP2* | up | 2 | 0.349 | 0 | 0.5 | 0.849 | 2.849 |
| 50 | *RNF123* | up | 2 | 8.00E-05 | 0 | 0.75 | 0.75008 | 2.75008 |
| 51 | *APOC1* | down | 2 | 0.21685 | 0 | 0.5 | 0.71685 | 2.71685 |
| 52 | *CEP57* | down | 2 | 0.072 | 0 | 0.5 | 0.572 | 2.572 |
| 53 | *PLPPR1* | down | 2 | 0.566 | 0 | 0 | 0.566 | 2.566 |
| 54 | *ADAM12* | up | 2 | 5E-07 | 0 | 0.5 | 0.5000005 | 2.5000005 |
| 55 | *ASCC3* | up | 2 | 1E-20 | 0 | 0 | 1E-20 | 2 |

**Fig. S1.**
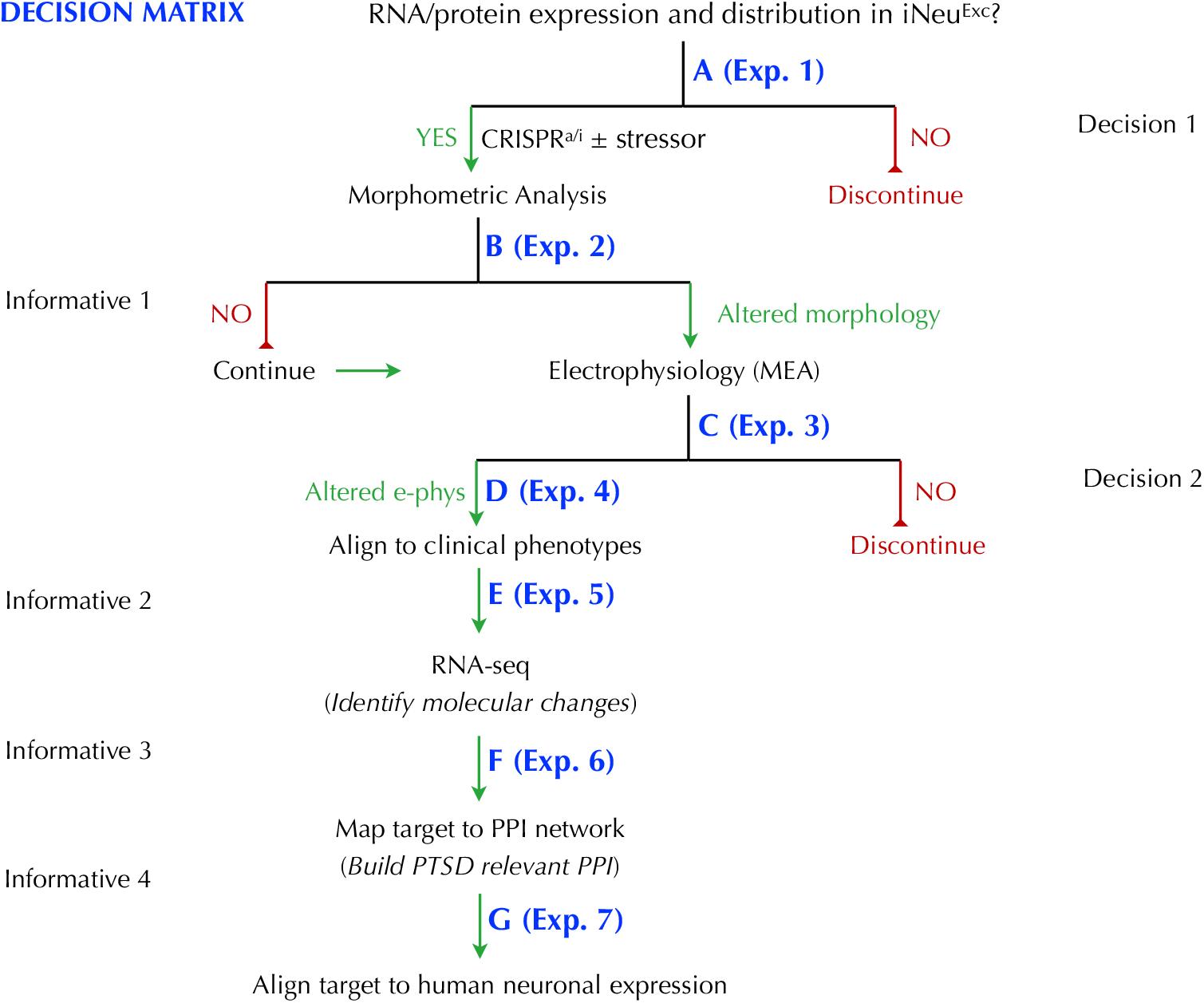
If transcript is expressed in iNeu^Exc^ capture pattern and conduct CRISPR^a/i^ modulation experiment followed by morphometric analysis; otherwise discontinue (**A**, Decision 1). Note: iNeu^Exc^ +/- CRISPR^a/i^ +/- stressor (oxytocin^**11**^ or hydrocortisone^**12**^), will be directly compared for decision matrix. Expression in iNeu^Exc^ is the first, but not terminal, decision point —> assess in iNeu^Inh^ (Part II). *If not expressed in either neuronal subtype, assessment of SNP-AG will be discontinued*. Morphometric analysis will next be done concomitant with neuronal marker labeling and although informative, it is not decision making (**B**). The next critical decision making step involves MEA: if electrophysiological (e-phys) properties are not altered as a result of CRISPR^a/i^ +/- stressor treatment then SNP-AG assessment is discontinued in iNeu^Exc^ (**C**, Decision 2). If e-phys is altered, capture changes and align expression/e-phys results to distinct known clinical phenotypes (**D**), which is potentially informative. Subsequently, conduct RNA-sequencing (RNA-seq) to identify changes in molecular signature of iNeu^Exc^ resulting from CRISPR^a/i^ modification of SNP-AG (**E**). Predicted molecular changes, concomitant with altered e-phys, are alterations in ion channel expression and synaptic-marker expression. Map SNP-AG to core PPI network to build understanding of protein relationships in the context of PTSD phenotypes (**F**). While expression of SNP-AG in human CNS has been ascertained using GTeX a follow-up to describe expression analysis in human neuronal subtypes and CNS tissue will be conducted (**G**).

**Fig. S2.**
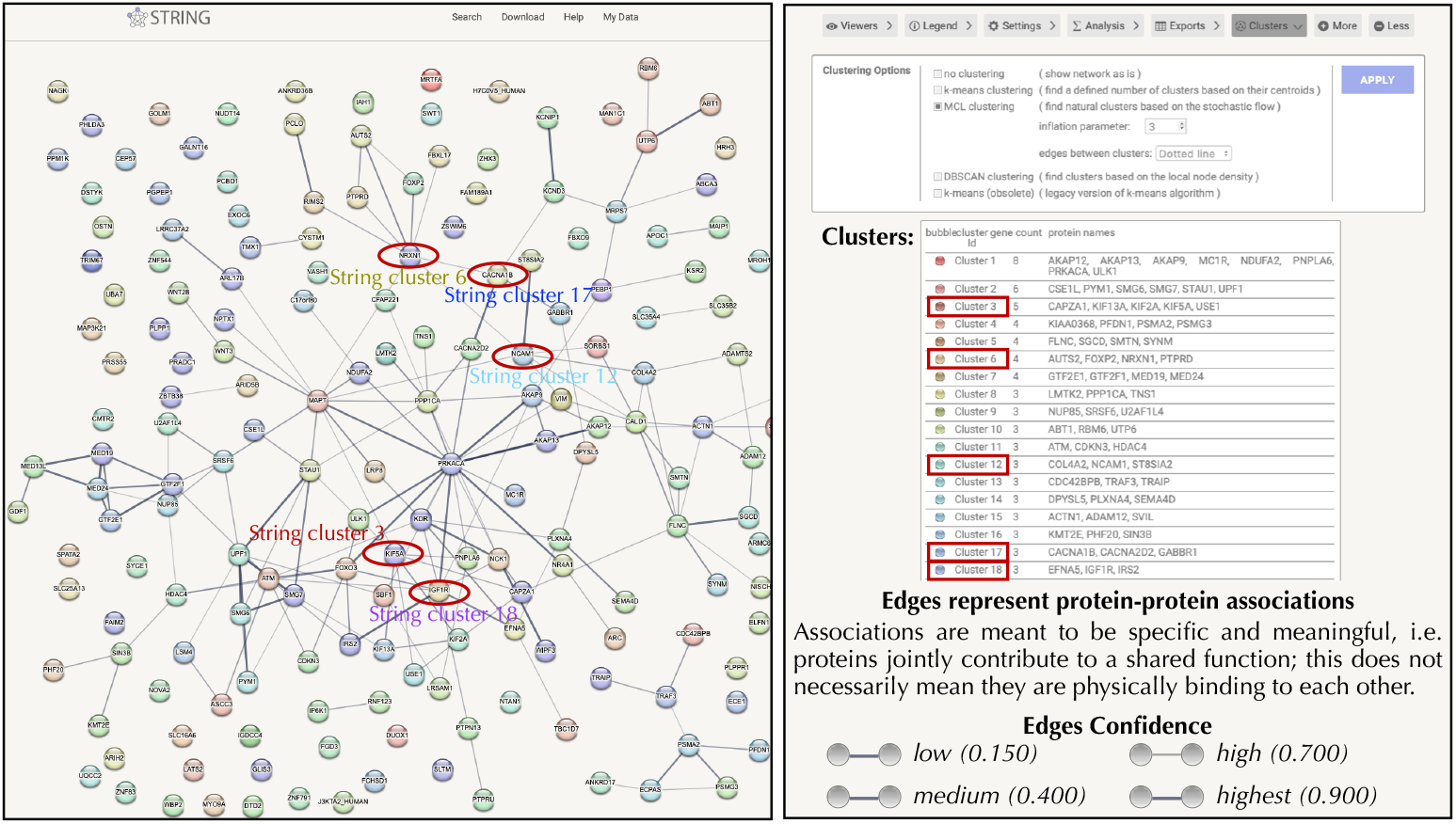
A PTSD PPI network was generated, based on the 177 Phase 1 list, by entering proteins into STRING using the multiple proteins search tool. Then, using STRING tools we interrogated the list using the <u>M</u>arkov <u>Cl</u>uster (MCL) Algorithm, which is a fast and scalable unsupervised cluster algorithm for networks based on simulation of (stochastic) flow in graphs (http://www.micans.org/mcl/). Nodes reflecting five clusters of interest are indicated by red ovals; Related edges are shown by red lines (**Left**). Details and the top 18 clusters/edges are shown in table format (**right**).

**Supplemental Table 6.** Top 20 Target Expression Summary. This table displays high-level information about the top 20 targets. “Type” refers to the protein class. Gene expression level (TPM) of each target is shown for frontal cortex (from GTEx-v8) and four brain cell types (excitatory neurons, inhibitory neurons, astrocytes, microglia; from Human Protein Atlas). Target names in blue (n=11) are supported by both transcriptomic and genomic evidence of association to PTSD.

| Target | Type | Frontal Cortex | Excitatory Neurons | Inhibitory Neurons | Astrocytes | Microglia |
| --- | --- | --- | --- | --- | --- | --- |
| CACNA1B | Ion Channel | 17.1 | 375.5 | 316.8 | 41.2 | 17.0 |
| IGF1R | Receptor (TRK) | 8.38 | 231.4 | 186.7 | 149.6 | 234.4 |
| HDAC4 | Enzyme (Deacetylase) | 8.7 | 114.0 | 81.4 | 58.8 | 36.7 |
| CACNA2D2 | Ion channel | 3.6 | 7.5 | 34.6 | 4.0 | 1.8 |
| KCND3 | Ion Channel | 23.7 | 102.1 | 84.6 | 17.3 | 0.9 |
| GABBR1 | Receptor (GPCR) | 123.8 | 56.7 | 65.2 | 142.7 | 18.8 |
| NDUFA2 | Enzyme (dehydrogenase) | 57.4 | 6.3 | 7.3 | 9.9 | 5.4 |
| PTPRD | Enzyme (phosphatase) | 22.9 | 4422.9 | 1105.1 | 188.9 | 81.4 |
| KIF5A | Enzyme (kinesin) | 839.3 | 27.5 | 20.8 | 2.8 | 0.0 |
| NRXN1 | Receptor (Transmembrane) | 18.2 | 3094.7 | 2015.5 | 3817.5 | 166.4 |
| MED13L | Scaffold | 6.8 | 301.0 | 307.4 | 288.3 | 408.0 |
| PCLO | Scaffold | 8.6 | 632.9 | 715.2 | 38.4 | 48.3 |
| NCAM1 | Cell adhesion molecule |  | 445.6 | 421.9 | 503.5 | 34.0 |
| FLNC | Attachment Protein | 3.9 | 0.2 | 1.0 | 0.6 | 0.0 |
| HRH3 | Receptor (GPCR) | 26.1 | 4.1 | 22.7 | 0.0 | 0.0 |
| KMT2E | Enzyme (methyltransferase) | 12.7 | 154.8 | 142.5 | 102.8 | 142.2 |
| FOXP2 | Transcription Factor | 1.0 | 179.9 | 6.1 | 5.3 | 275.5 |
| KIF2A | Enzyme (kinesin) | 14.7 | 54.7 | 61.8 | 51.7 | 25.0 |
| KDR | Receptor (RTK) | 2.3 | 0.1 | 1.3 | 0 | 7.2 |
| TRAF3 | Enzyme (ligase) | 9.2 | 104.7 | 60.5 | 51.7 | 22.4 |

**Supplemental Table 7.**
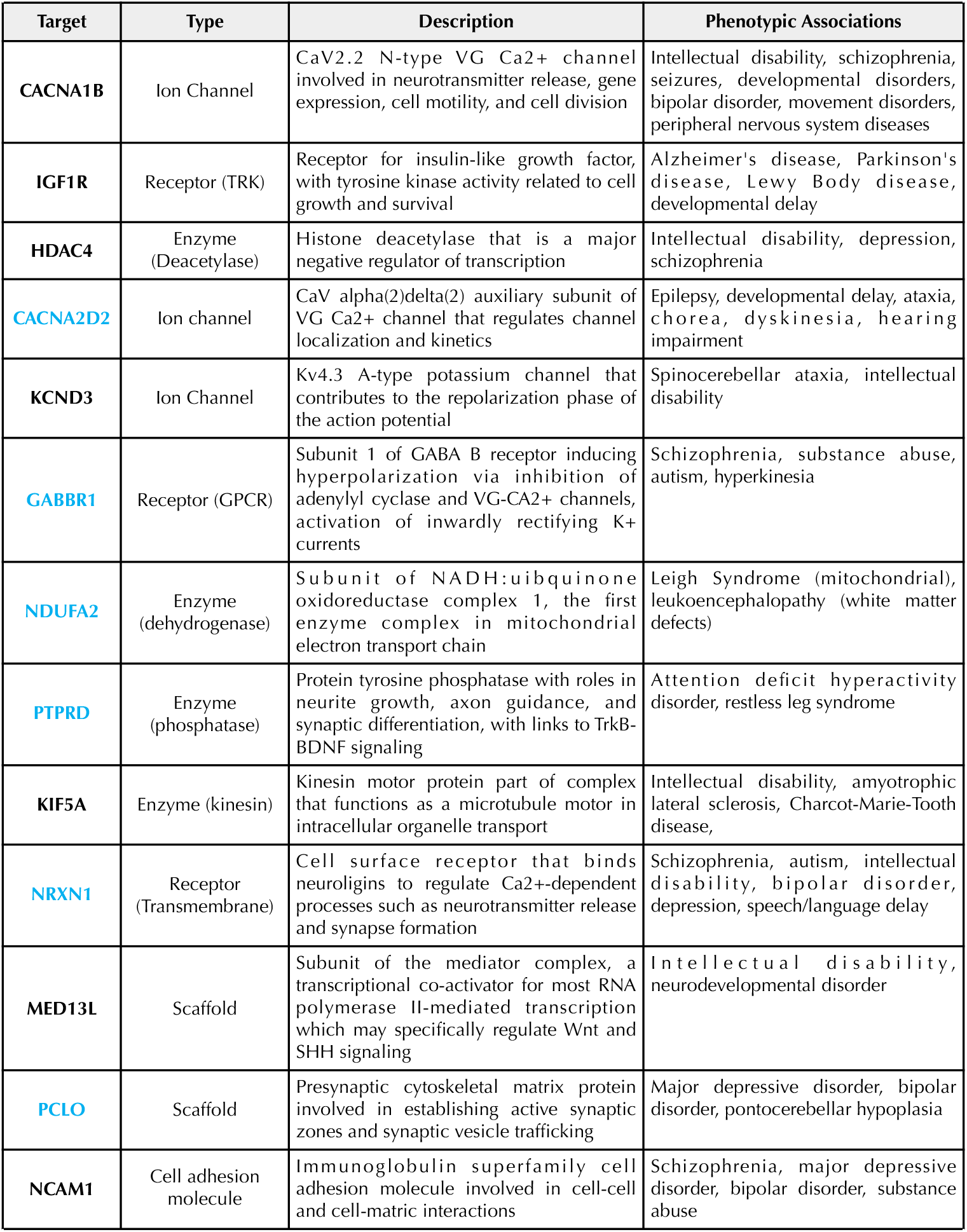
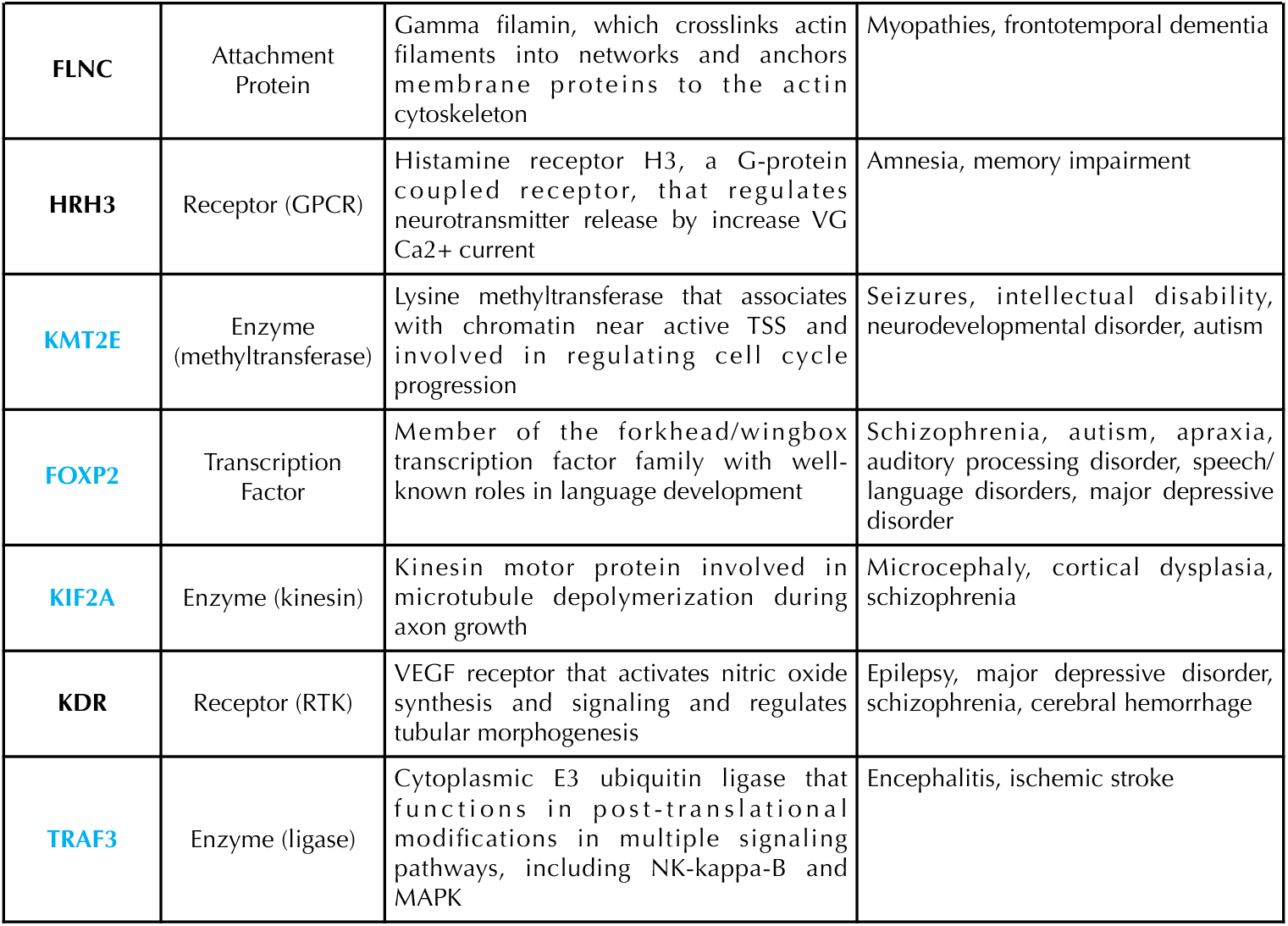
Top Twenty Target Biology Summary.

| Target | Type | Description | Phenotypic Associations |
| --- | --- | --- | --- |
| **CACNA1B** | Ion Channel | CaV2.2 N-type VG Ca<sup>2+</sup> channel involved in neurotransmitter release, gene expression, cell motility, and cell division | Intellectual disability, schizophrenia, seizures, developmental disorders, bipolar disorder, movement disorders, peripheral nervous system diseases |
| **IGF1R** | Receptor (TRK) | Receptor for insulin-like growth factor, with tyrosine kinase activity related to cell growth and survival | Alzheimer's disease, Parkinson's disease, Lewy Body disease, developmental delay |
| **HDAC4** | Enzyme (Deacetylase) | Histone deacetylase that is a major negative regulator of transcription | Intellectual disability, depression, schizophrenia |
| **CACNA2D2** | Ion channel | CaV alpha(2)delta(2) auxiliary subunit of VG Ca<sup>2+</sup> channel that regulates channel localization and kinetics | Epilepsy, developmental delay, ataxia, chorea, dyskinesia, hearing impairment |
| **KCND3** | Ion Channel | Kv4.3 A-type potassium channel that contributes to the repolarization phase of the action potential | Spinocerebellar ataxia, intellectual disability |
| **GABBR1** | Receptor (GPCR) | Subunit 1 of GABA B receptor inducing hyperpolarization via inhibition of adenylyl cyclase and VG-CA<sup>2+</sup> channels, activation of inwardly rectifying K<sup>+</sup> currents | Schizophrenia, substance abuse, autism, hyperkinesia |
| **NDUFA2** | Enzyme (dehydrogenase) | Subunit of NADH:ubiquinone oxidoreductase complex 1, the first enzyme complex in mitochondrial electron transport chain | Leigh Syndrome (mitochondrial), leukoencephalopathy (white matter defects) |
| **PTPRD** | Enzyme (phosphatase) | Protein tyrosine phosphatase with roles in neurite growth, axon guidance, and synaptic differentiation, with links to TrkB-BDNF signaling | Attention deficit hyperactivity disorder, restless leg syndrome |
| **KIF5A** | Enzyme (kinesin) | Kinesin motor protein part of complex that functions as a microtubule motor in intracellular organelle transport | Intellectual disability, amyotrophic lateral sclerosis, Charcot-Marie-Tooth disease, |
| **NRXN1** | Receptor (Transmembrane) | Cell surface receptor that binds neuroligins to regulate Ca<sup>2+</sup>-dependent processes such as neurotransmitter release and synapse formation | Schizophrenia, autism, intellectual disability, bipolar disorder, depression, speech/language delay |
| **MED13L** | Scaffold | Subunit of the mediator complex, a transcriptional co-activator for most RNA polymerase II-mediated transcription which may specifically regulate Wnt and SHH signaling | Intellectual disability, neurodevelopmental disorder |
| **PCLO** | Scaffold | Presynaptic cytoskeletal matrix protein involved in establishing active synaptic zones and synaptic vesicle trafficking | Major depressive disorder, bipolar disorder, pontocerebellar hypoplasia |
| **NCAM1** | Cell adhesion molecule | Immunoglobulin superfamily cell adhesion molecule involved in cell-cell and cell-matrix interactions | Schizophrenia, major depressive disorder, bipolar disorder, substance abuse |
| **FLNC** | Attachment Protein | Gamma filamin, which crosslinks actin filaments into networks and anchors membrane proteins to the actin cytoskeleton | Myopathies, frontotemporal dementia |
| **HRH3** | Receptor (GPCR) | Histamine receptor H3, a G-protein coupled receptor, that regulates neurotransmitter release by increase VG Ca2+ current | Amnesia, memory impairment |
| **KMT2E** | Enzyme (methyltransferase) | Lysine methyltransferase that associates with chromatin near active TSS and involved in regulating cell cycle progression | Seizures, intellectual disability, neurodevelopmental disorder, autism |
| **FOXP2** | Transcription Factor | Member of the forkhead/wingbox transcription factor family with well-known roles in language development | Schizophrenia, autism, apraxia, auditory processing disorder, speech/language disorders, major depressive disorder |
| **KIF2A** | Enzyme (kinesin) | Kinesin motor protein involved in microtubule depolymerization during axon growth | Microcephaly, cortical dysplasia, schizophrenia |
| **KDR** | Receptor (RTK) | VEGF receptor that activates nitric oxide synthesis and signaling and regulates tubular morphogenesis | Epilepsy, major depressive disorder, schizophrenia, cerebral hemorrhage |
| **TRAF3** | Enzyme (ligase) | Cytoplasmic E3 ubiquitin ligase that functions in post-translational modifications in multiple signaling pathways, including NK-kappa-B and MAPK | Encephalitis, ischemic stroke |

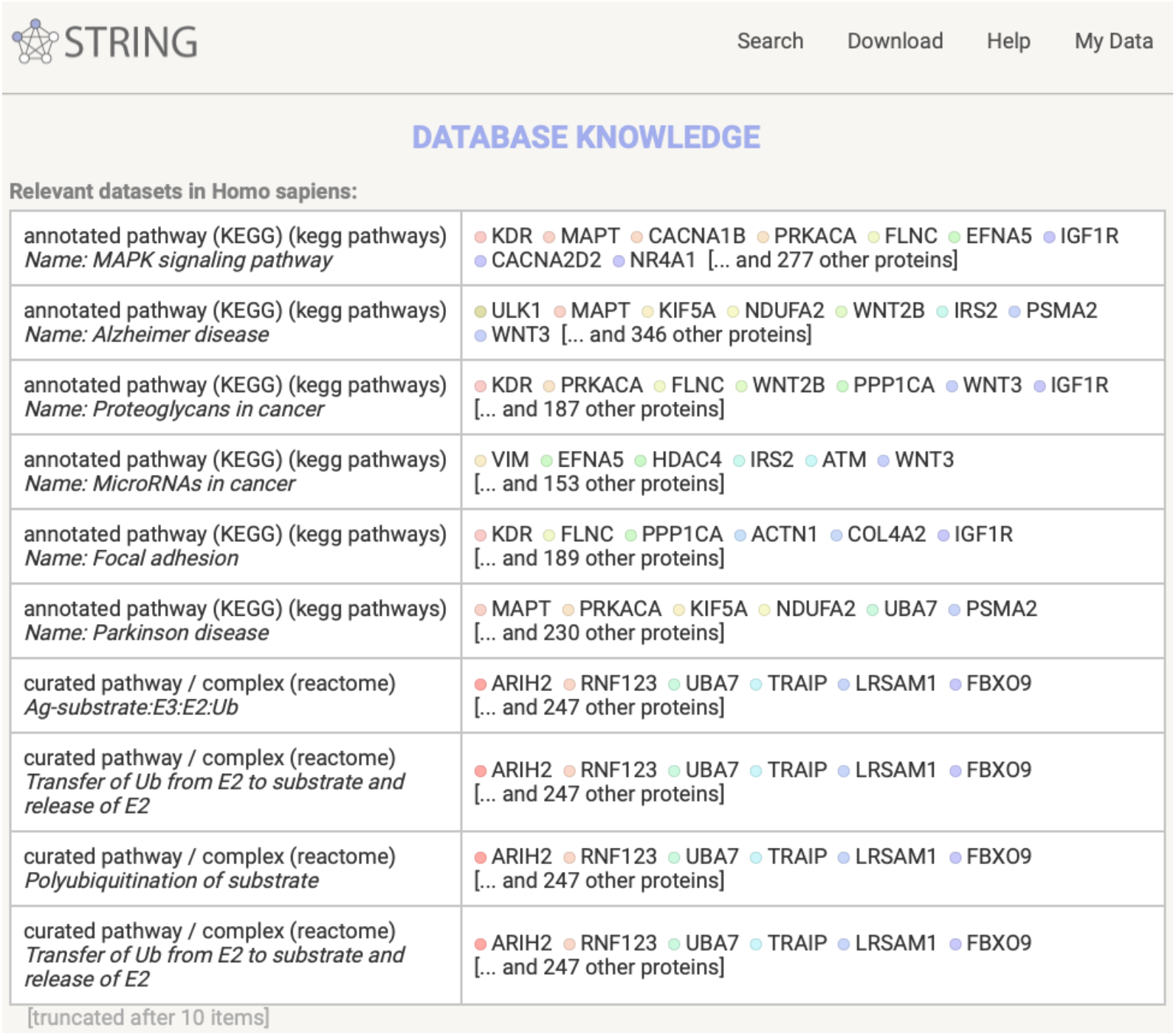

### DisGeNET

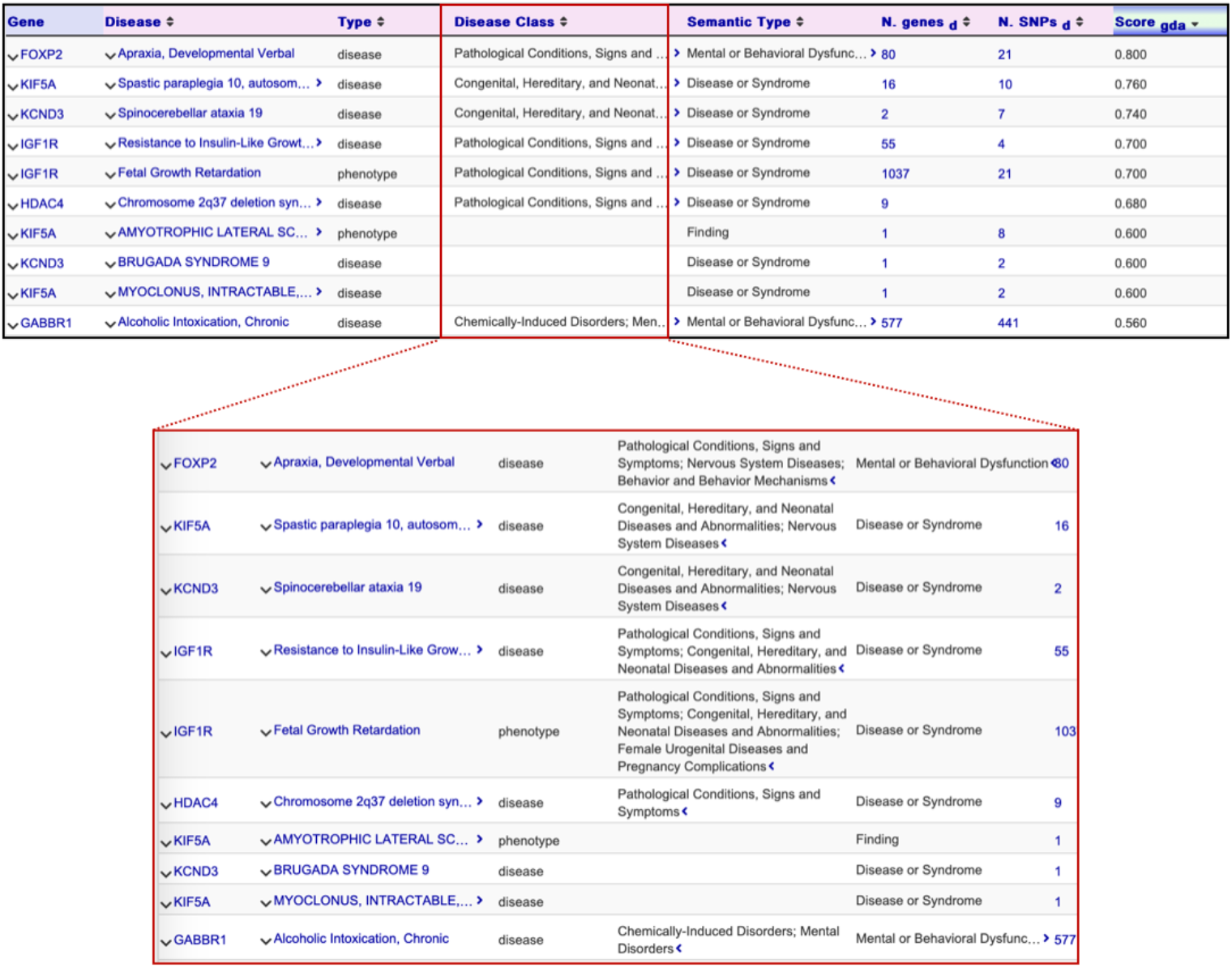

**Supplemental Table 8.** PTSD TAP Putative Therapeutics. From our top 20 PTSD targets, five (25%, at the time of assessment) had therapeutic molecules at various stages of clinical development for CNS indications, which suggests that they could be used to validate utility for PTSD cell based phenotypes as determined by our Decision Matrix (**Fig. S1**)

| Putative Target | Disease | Drug | Mechanism | Phase | Validity |
| --- | --- | --- | --- | --- | --- |
| CACNA1B | Epilepsy | Levetiracetam | N-Type Calcium Channel (Ca(v) 2.2) Blockers | Launched | Validated |
| CACNA1B | Epilepsy | Verapamil | Cytochrome P450 CYP1A2 Inhibitors | Phase II | Exploratory |
| CACNA1B | Epilepsy | Verapamil | Cytochrome P450 CYP2C8 Inhibitors | Phase II | Exploratory |
| CACNA1B | Epilepsy | Safinamide mesilate | Calcium Channel Modulators | Phase II | Exploratory |
| CACNA1B | Epilepsy | dl-cis-ML-1018 | Calcium Channel Blockers | Biological Testing | Exploratory |
| GABBR1 | Epilepsy | NCS-399 | GABA type B receptor subunit 1 Modulator | Biological Testing | Exploratory |
| HDAC4 | Epilepsy | Valproate semisodium | Epoxide Hydrolase Inhibitors | Launched | Validated |
| HDAC4 | Neurodegeneration | EHT-0205 | Histone Deacetylase (HDAC) Inhibitors | Preclinical | Candidate |
| IGF1R | Neurodegeneration | NG-56 | Signal Transduction Modulators | Biological Testing | Exploratory |
| IGF1R | Neurodegeneration | NG-35 | Tyrosine Kinase Inhibitors | Biological Testing | Exploratory |
| IGF1R | Neurodegeneration | NG-35 | Signal Transduction Modulators | Biological Testing | Exploratory |
| KDR | Epilepsy | SG-00529 | mTOR Complex 1 (mTORC1) Inhibitors | Preclinical | Validated |
| KDR | Epilepsy | SG-00529 | mTOR Complex 2 (mTORC2) Inhibitors | Preclinical | Validated |
| Source: Legacy version of the Integrity Discovery Database (https://discoverydata.com) |  |  |  |  |  |

## Notes

https://drive.google.com/drive/folders/1FaG0E_5ConmyUvuuV_LznwsHQjoxpoJs?usp=sharing

### Competing Interest Statement

The authors have declared no competing interest.

